# Extinctions have strongly reduced the mammalian consumption of primary productivity

**DOI:** 10.1101/2020.10.15.341297

**Authors:** Rasmus Østergaard Pedersen, Søren Faurby, Jens-Christian Svenning

## Abstract

Herbivorous mammals are important for natural ecosystems even today, but how much stronger would there effects be without human-linked extinctions and extirpations? The ranges of many mammal species have contracted and numerous species have gone extinct due to human pressures, so herbivore impacts in even seemingly natural ecosystems likely deviate from their pre-anthropogenic state. However, such effects remain poorly understood and often unrecognized. To address this issue, we here quantified and mapped plant consumption by all terrestrial mammals in natural areas based on both current and estimated natural ranges. We then compared the estimated consumption rates to current plant net primary productivity, and summarised the results for global ecosystem types both broadly and in the wildest remaining natural areas around the world (the Last of the Wild). We found that wild mammals consume 7.3% (95% interquantile range: 0.85% - 26%) of net primary productivity in current natural areas, and that this would be much higher in the absence of extinctions and extirpations, namely 13% (95% interquantile range: 1.7% - 40%), i.e., a >50% higher consumption rate. Marked human-linked declines in herbivory were seen even in the wildest remaining natural areas, where mammals now consume a mean of 9% (95% interquantile range: 2.2% - 26%) of plant primary productivity, which is only 60% of no-extinction level. Our results show that mammalian herbivores naturally play an important part in ecosystems at a global scale, but that this effect has been strongly reduced by extinctions and extirpations.

## Introduction

There is wide evidence that large herbivorous mammals can strongly shape vegetation^1–3^. Nevertheless, the general importance of such effects is poorly understood, with several studies reporting that mammalian herbivores consume a surprisingly small amount of the net primary production^4^. At the same time, large herbivore assemblages have been strong affected by human activities^5,6^, meaning that we have poor understanding of natural levels of herbivory and their vegetation effects. During the last 100,000 years, modern humans have spread across the whole world and with their arrival a large part of the megafauna has disappeared^7,8^. Only parts of Africa and small parts of Asia have retained megafauna assemblages comparable to those which once roamed the whole planet^6^. When these megafauna have been released from human pressures in protected conservancies, they have a large effect on ecosystem and vegetation structure^5,9^.

The ecosystem impact of a species tends to scale with body mass making the changes in ecosystems following megafauna extinctions larger than the changes in mammal diversity. Large animals can both have major direct structural effects (e.g. when elephants knock over trees^10^), but are also disproportionally important in indirect ways like overall vegetation consumption since a species’ energy requirements over a given area tends to increase with body mass^11,12^. The exact consequences of the prehistoric and historic megafauna range contractions extinctions on vegetation structure and ecosystem function is poorly known, albeit a rising number of studies point to widespread major effects: South American savannas would have been much more open like the African savannas^13^. Beetle assemblages from Great Britain indicate both a larger proportion of dung and more open and diverse mixture of vegetation cover in the Last Interglacial than Early Holocene^1^. A study from Queensland showed that megafauna extinction and subsequent increased wildfires led to the shift from mixed savanna including fire-sensitive trees to fire-tolerant sclerophyll vegetation^14^, and strong vegetation changes have also been coupled to megafauna losses in northeastern North America^15^. However, the large and cascading effects of large herbivores are also evident from the remaining species^5^, and the few areas with well-developed wild herbivore faunas that still exist (e.g. extirpation of the large herbivores in Mozambique led to an expansion in an invasive species and reintroductions took the invasive back to pre-extirpation occurence^16^, re-establishment of bison numbers in Yellowstone National Park are limiting the woody plant communities^17^, exclosure experiments in temperate forests have shown that saplings have a hard time escaping herbivores both under closed forest canopy and in large gaps^18^, and long-term elephant use decreases vegetation height and increases vegetation height variablity^19^). It is obvious that large herbivores have a unique importance in ecosystem function and are irreplaceable by smaller species and this has implications for nature restoration^5,20^. Even though the extinct Late Pleistocene terrestrial megafauna only made up a small fraction of the global species diversity of mammals (3.9%), their effect on vegetation seem likely to have been substantial^3^.

A review of consumption studies on current mammalian terrestrial fauna have shown them to have a highly variable, but often quite small effect on total net primary productivity (NPP), consuming only a median of 2%, although varying from <1% to 29%^4^. A few macroecological studies have made rough estimates of the effect of extinctions on the consumption patterns and have estimated that the extinction of megafauna have decreased mammalian consumption by 2.2% - 5.3% of NPP^21^. Earlier studies, however, use broad allometric scaling equations for density and consumption estimates based on a limited number of data points, even though we know that different functional and taxonomic groups can scale widely different^12^. Here we use taxonomically wide datasets and use phylogenetic models to estimate individual species density and metabolic rates. So even though the impact of megafauna seems small, there has clearly been a significant drop in impact as a consequence of the megafauna extinctions. Only few studies have tried to address and understand the importance of mammals for regulating vegetation growth at a global scale. How important would they be, if there had been no megafauna extinctions and extirpations in the late Quaternary?

Here, we assess the potential global impact of herbivory by current and present-natural mammalian fauna (i.e., under current climatic conditions but in the absence of anthropogenic late-Quaternary extinctions and extirpations). We have assembled and combined data on ranges, metabolic demands, population densities, and diets for all mammals extant throughout the last 130,000 years. Putting this together enabled us to estimate the overall consumption of the planet’s vegetation production by mammals.

We compare this to estimates of current plant productivity to estimate how large effects mammals with and without megafauna losses have on the ecosystems. Even though only a small part of the mammalian fauna has gone extinct, we (1) expect that the extinctions have had a strong impact on vegetation consumption. We compare the losses across realms and biomes, to assess how consistent the patterns are geographically and ecologically. We (2) expect to see a consistent pattern geographically, with least impact in the Afrotropics which still has the most extant megafauna. Forested areas have been least impacted by humans and we could therefore (3) expect a smaller impact in those ecosystems. The wildest places left on earth, have not been immune to megafauna extinctions and we therefore (4) expect to see the same patterns even there.

## Results and discussion

### Wild mammal biomass

We estimated wild mammal biomass across the globe, assuming total coverage and full natural density within each 96 km × 96 km grid cell. This was done for current-day non-introduced ranges for all extant mammal species, as well as for present-natural ranges of all late-Quaternary mammal species, i.e., without any late-Quaternary extinctions or any human-driven range modifications.

We estimate present-natural terrestrial mammal carbon stock (outside deserts) to be 0.32 PgC (95% interquantile range: 0.047 PgC −2.9 PgC). This is substantially higher than previous estimates. A study^22^ of mammal biomass through time estimated the pre-extinction biomass to be around 0.03 PgCs, but using different methods and cruder ranges. A study of current global carbon in livestock estimate 0.1 PgC in livestock and 0.003 PgC in wild mammal populations^23^. So current-day mammal biomass including livestock is comparable to the potential-natural carbon stock in wild animals, just with a near-complete shift from wild to domestic animals. We note that these numbers only concern standing biomass and not biomass flux, a faster turn-over rate in domestic animals would mean a much higher consumption per kg farmed biomass relative to wild biomass.

Mapping total mammal biomass, we get comparable estimates to what others have found before, based on empirical local animal counts^24^. Fløjgaard et al.^24^ report a bimodal distribution of empirical large-herbivore biomass in Europe with the high numbers, from rewilding sites, in line with our estimates. Comparing to the empirical values from their study based on animal counts we find that our theoretical estimates are broadly in agreement, but for most areas in the Afrotropics tend to be higher than those observed empirically (Fig S6). The latter can likely be attributed to reduced megafauna densities due to hunting and other human pressures^25,26^. The top-level empirical from the Afrotropics, such as from the Maasai Mara, Kenya with its nearly intact megafauna, are close to ours, despite even these areas being affected by anthropogenic megafauna declines^27^, further supporting that our estimates are reasonable equilibrium estimates in the absence of human pressures.

### Current vegetation consumption by wild mammals

When we estimate current plant consumption by wild mammals, we note that this is meant for *natural areas*. We state this to highlight that most places are under anthropogenic use, and that our numbers are applicable where wild animal species occur within the cell at their natural densities. Our estimates suggest that current plant consumption by wild mammals in natural areas varies greatly across the globe, with the highest consumption in western North America, central and eastern Africa, as well as a large spike in central Asia (Fig 2). This pattern largely follows the current diversity of large mammals across the globe^6^, with the most diverse areas having the highest consumption. We compared how our estimates of mammal plant consumption compares to net primary productivity (NPP), and found the median consumption by wild mammal populations was only around 7.3% (mean 9.1%) (95% interquantile range: 0.85% - 26%) (Fig 2–4). Areas like tropical South America and coastal Australia have very few extant large mammal species but high plant productivity, therefore almost none of the NPP is currently being consumed by mammals in these ecosystems. Our estimate of current consumption as a percentage of NPP fits well with empirical studies: A review on NPP consumption in modern terrestrial ecosystems found that across 21 studies, animals eat 0.24% - 29% of the total production with a median of 1.9%^4^.

In a study^28^ on one of the most megafauna rich ecosystems today, the Serengeti National Park, Tanzania, large-herbivore consumption at the grazed sites was found to be median 163 Mg Carbon / km^2^ / year (range 36-462, n = 28, extracted from figures in the paper), i.e., 15% of our estimate of total NPP for the area (1115 Mg Carbon / km^2^ / year). Our estimate from the same area is that that 112 Mg Carbon / km^2^ /year is currently consumed, i.e., 10% of total NPP. Hence, our estimate is close the empirical estimates in intact ecosystems.

### Differences between current and present-natural vegetation consumption

Global plant consumption by mammals in natural areas is very different now compared to what it would have been today had a large part of the mammalian megafauna not been lost (Fig 1), with especially large differences in parts of the Americas and Australia, where >50% of the present-natural mammalian consumption is missing today (Fig. 1 and 3), i.e., considering only native species and ignoring the impacts of introduced species^29^. Wild areas in Africa and Asia where more of the megafauna has been preserved are much closer to present-natural consumption. The global median fraction of NPP consumed for present-natural distributions is 13% (mean 15%) (95% interquantile range: 1.7% - 40%) of NPP, i.e., 1.8 times the current level (Fig 2–4).

**Fig 1:**
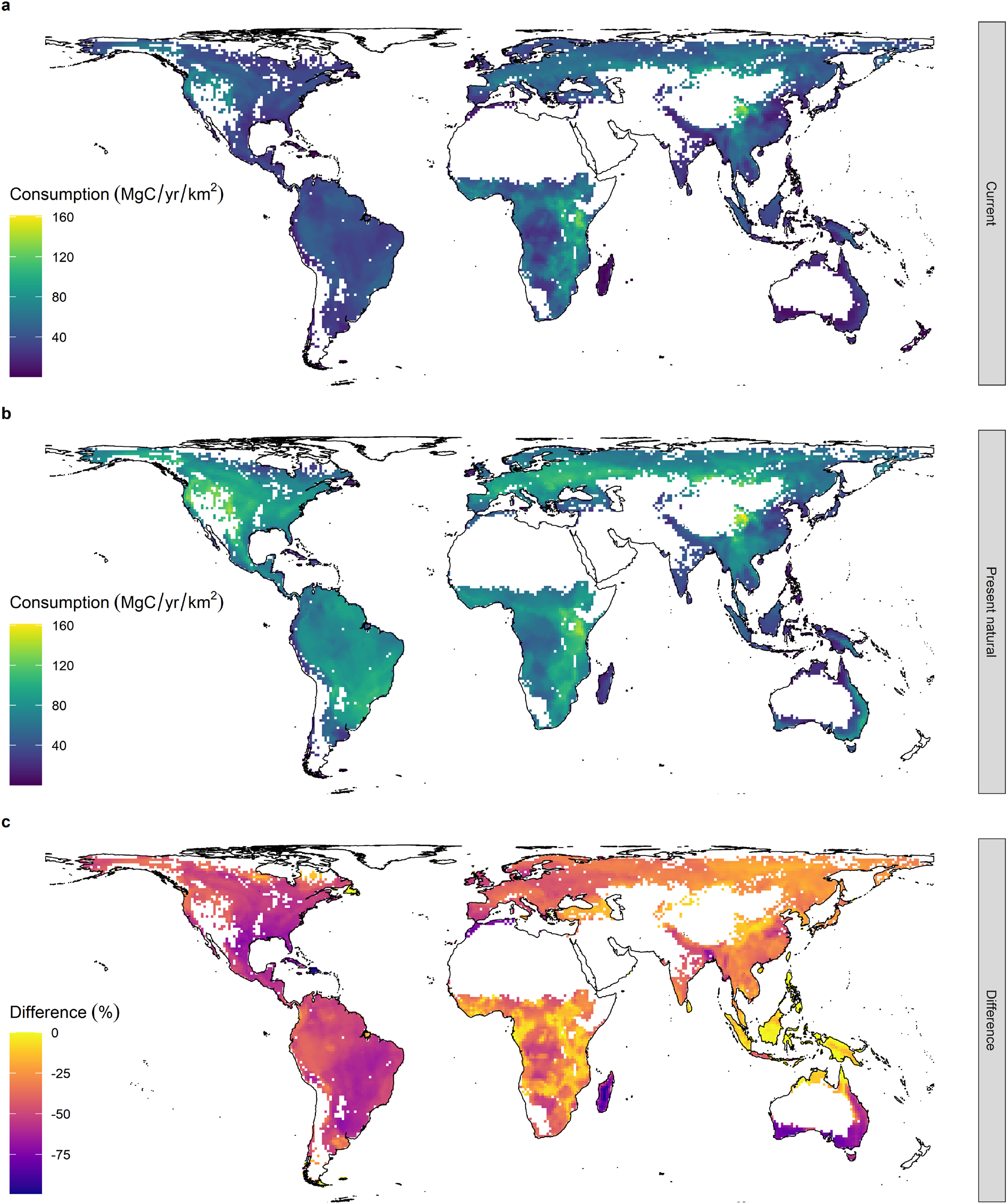
Mammal consumption of plant carbon globally. a) Average terrestrial mammal consumption by current mammals (current ranges of extant species). b) shows present natural consumption (potential present day ranges with no human presence). c) shows the percentage lower consumption of current ranges compared to present natural Blank pixels represents areas excluded from the analyses due to very low NPP or high variability in NPP. (See Fig S1 for included low NPP regions.)

**Fig 2:**
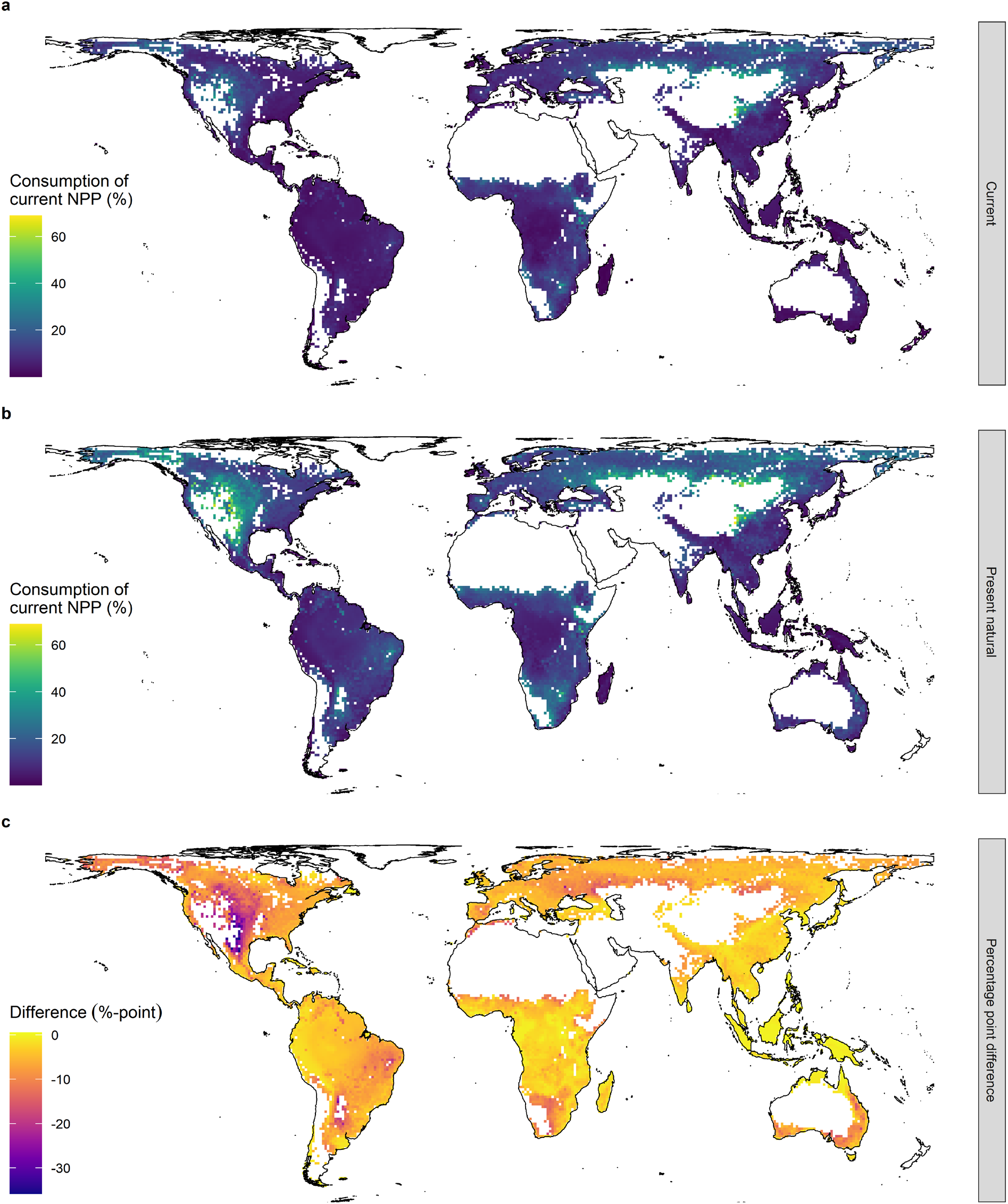
Estimated mammal consumption of net primary productivity (NPP) globally. a) Consumption by current mammals (current ranges of extant species). b) Present natural consumption (potential present day ranges with no human presence). c) Percentage-point consumption difference between present natural and current consumption. Blank pixels represents areas excluded from the analyses due to very low NPP or high variability in NPP. (See Fig S2 for included low NPP regions.)

**Fig 3:**
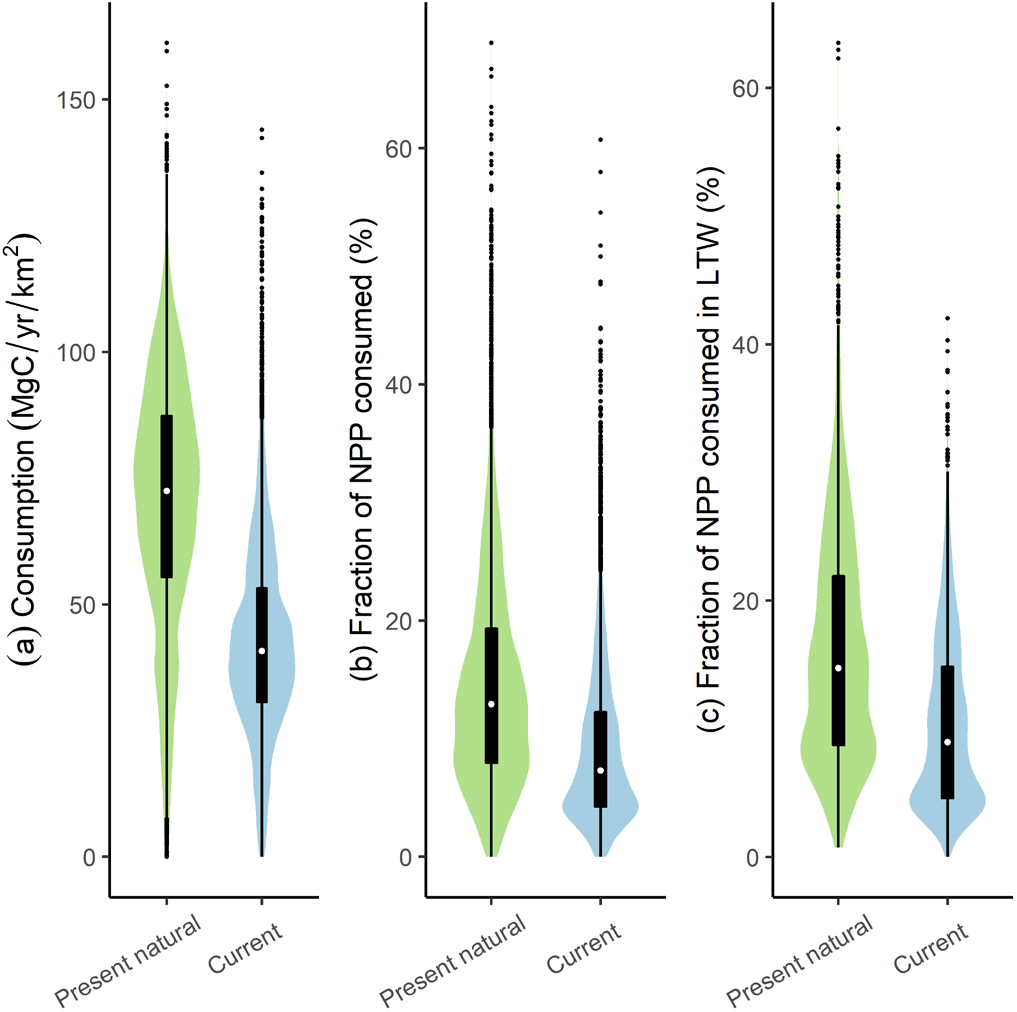
Global consumption summary. Boxplot with underlying violins with kernel density scaled to width. **a)** Total consumption of carbon. **b)** Fraction of net primary productivity (NPP) consumed. **c)** Fraction of net primary productivity (NPP) consumed in the areas designated as ‘last of the wild’^37,50^. (See Fig S3 for included low NPP regions.)

**Fig 4:**
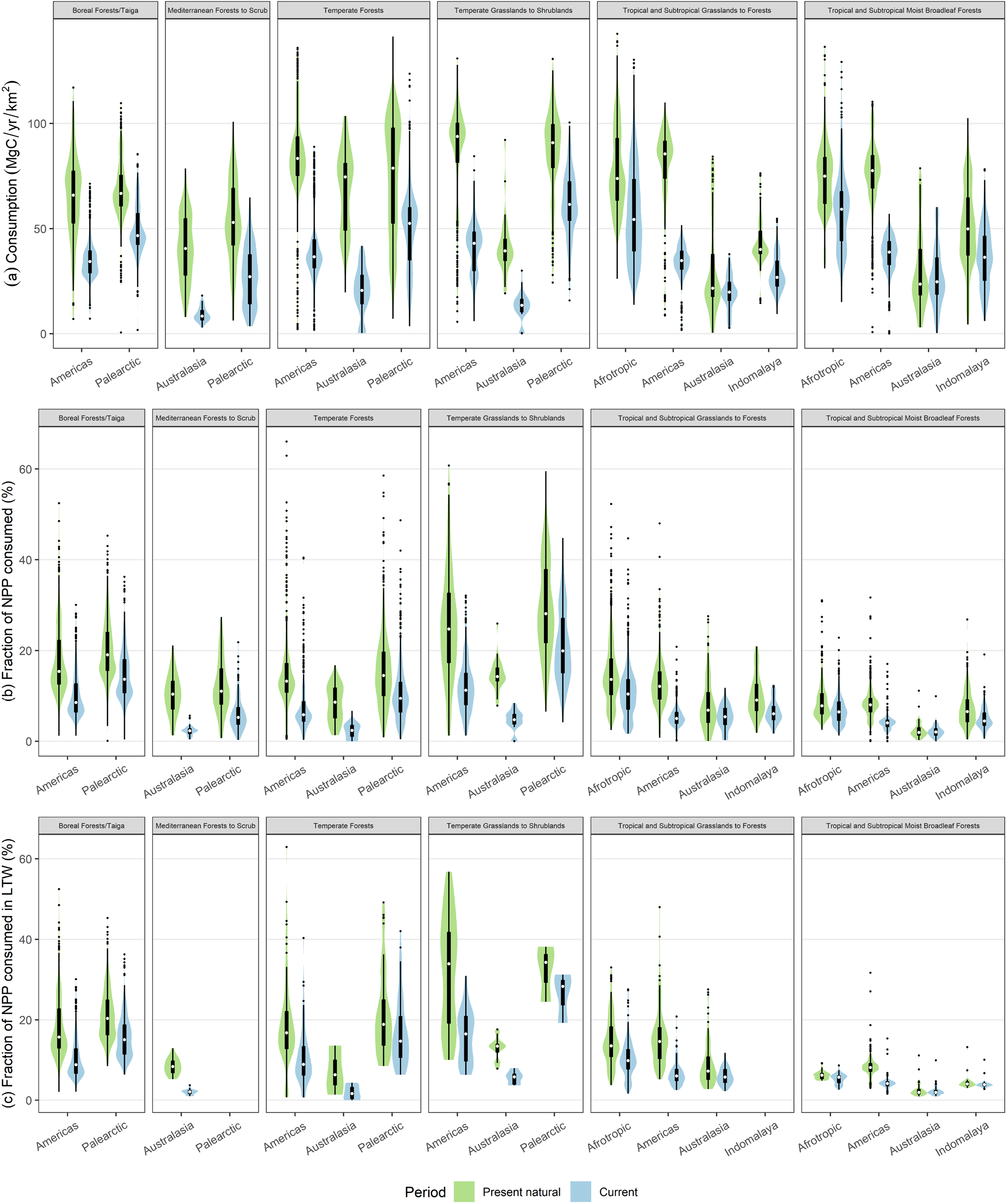
Consumption summary for large ecological units based on realms and biomes^51^. Boxplot with underlying violins with kernel density scaled to width. Neotropics and Nearctic are merged to the Americas and Madagascar is excluded from the Afrotropics for biome modifications see Fig S5. **a)** Total consumption of carbon. **b)** Fraction of net primary productivity (NPP) consumed. **c)** Fraction of net primary productivity (NPP) consumed in the areas designated as ‘last of the wild’^37,50^. (See Fig S4 for included low NPP regions.)

As increased mammal density may sometimes increase NPP, our use of current NPP might in some areas underestimate of present-natural NPP. Several insect studies have shown increased productivity in the presence of increased herbivory and viceversa^30,31^. Mammals have also been shown to increase productivity. Arctic tundra ecosystems are nutrient-limited and mammals increase nutrient turnover and in turn increase the productivity of grasslands – which they likely did so much more in the past^32^. Further, both in Africa and South America grazing lawns are being kept by large herbivores, again increasing producticvity^33^, and a grazing study in Serengeti, showed that cutting and grazing stimulated productivity^28^. Hence, in such situations our estimated present-natural proportional consumption rates may be overestimates.

Only 7.4% of all terrestrial mammal species have gone extinct since the Late Pleistocene, but a massive 54% of terrestrial megafauna mammal species (≥45 kg body mass) have been lost across the same period^34,35^. Our results show that this have strongly changed NPP consumption levels. The current mammal fauna accounts for an average of 9.1% of plant consumption in natural areas, with megafauna accounting for 13% hereof. In the absence of extinctions and extirpations, the present-natural fauna would consume an average of 15% of plant consumption in natural areas, with megafauna accounting for a 45% hereof, i.e., megafauna loss accounts for 93% of the overall decrease in NPP consumption. As discussed earlier, there are current and paleoecological examples of how large herbivores can be very important for the environment and how their loss may cause large vegetation shifts across the world’s biomes^3^. This is reaffirmed in our consumption estimates with a 44% reduction in NPP consumption (13% to 7.3%).

Some regions are particular affected. For example, Australia’s native megafauna diversity is extremely impoverished compared to the present-natural diversity. Out of the 33 late-Quaternary species of strict herbivorous (≥ 90% plant diet) marsupials > 20 kg, only 8 are left (21 kg - 31 kg and one of 46 kg) of whom one is classified as threatened by IUCN (International Union for Conservation of Nature). While the median body size of the extinct fauna was 131 kg, and the heaviest 2700 kg^35,36^. This causes the extreme drops in consumption we see in the temperate grasslands, shrublands and forests of Australasia (Fig. 4), though some of this effect might be mitigated by introduced species that were not included in this study^29^.

### Reduced herbivore effects even in natural areas

In the areas with the lowest human footprint (‘Last of the Wild’^37^), the median fraction of NPP consumed is 9.0% (95% interquantile range: 2.2% - 26%) and 15% (95% interquantile range: 3.4% - 40%) for current and present-natural ranges, respectively (Fig. 3c). It has previously been highlighted that there is poor overlap between human footprint and faunistic intactness^38^. Our finding of strongly reduced large-herbivore consumption in the Last of the Wild areas shows that many of these apparently low-impact areas are not only substantially modified in their species composition, but also in their functional ecology.

### Perspective

The ecological implications of our findings are complex and will depend both on biomes and the ecological characteristics of the herbivores that has been lost. The consequences of reduced large mammal herbivory for woody vegetation and fire risk for instance depends of the relative dominance of browser vs grazers. The loss of many large browsers in the Americas could have reduced the fire risk, since browsers create woody debris and promote semi-open grasslands with flammable grasses^39^. On the other hand, we see in our model, that the full Pleistocene community would have consumed a much larger fraction of the plant productivity likely leading to much lower fuel loads, which should have decreased fire severity^40^. A dramatic loss of consumption of the plant productivity would also lead to a promotion of competitive plant species, and less seed dispersal^415,39^, potentially leading to reduced plant diversity (either at the landscape scale, due to vegetation composition shift^42^ or locally due to decreased connectivity^43^). Several studies have found that the extinction of megafauna have led to ecosystem impacts such as loss of open mosaic vegetation^1,17^ and increased fire^44^. The extinction of arctic megafauna likely led to an increase in *Betula* cover, with potential effects on global climate^32,45^. As an example changes in fire regimes a study in Australia found that the extinction of extinction megafauna preceded an increased fire regime which again preceded a vegetation shift from mixed savanna with both rainforest and sclerophyll trees to purely sclerophyll vegetation^14^. A review of current elephant impact found that while elephants decreases tree abundance, they do not affect their diversity and increases herb diversity, and found no consistent cascading effects on either abundance or diversity other animals^46^.

Whenever we look to seemingly pristine ecosystems^37^, we have to keep in mind that even those ecosystems are affected by human-linked ecological changes. With a large proportion of the terrestrial megafauna extinct or extirpated across the globe^6^, a large part of the mammal function has been lost^47,48^. As this study shows, even the “least impacted” environments on the planet, have also experienced a large reduction in mammal vegetation consumption. The mammal communities have been impoverished to such an extent that introduced species make up an important part of their functional space^29^. These extinctions must have had an effect on the ecosystems: Decreased mammal consumption leading to a variety of effects including, but not limited to on changes in community composition and structure^1,13,16,17,19^ or changes in fire frequency and intensity^14,15,39^. Our work cannot with certainty identify how ecosystems would have functioned prior to the late-Quaternary megafauna losses, but highlights that even remote ecosystems are likely fundamentally changed relative to a pre-extinction baseline^21^. The world we live in is drastically different from one without human impacts even in seemingly pristine landscapes. In consequence, our findings also have strong implications for ecosystem restoration, notably trophic rewilding^49^. Notably, they highlight that it is important to consider the strong down-sizing of large-herbivore assemblages since the Late Pleistocene and associated reduction in NPP consumption rates in efforts to restore self-sustaining ecosystems, e.g., via active megafauna restoration^49^.

## Methods

The goal of this paper was to estimate the total consumption by wild mammals in a non human-dominated world. To do this we estimated densities and energetic needs for all mammals and combined this with previously gathered information on diet and current and natural range size of all mammals. To estimate the total impact of consumption by mammals globally and to what degree this must be affected by the human impacts, we needed information on species distributions with and without human presence impacts, population densities, energetic needs, and diet.

### Consumption

#### Taxonomic scope

We followed the mammalian species list in PHYLACINE^34,35^ (v.1.2.1) which follows the IUCN Red List^52^ for extant species. This list was filtered to include only terrestrial not primarily marine species i.e. excluding bats, whales, pinnipeds and sea cows, and three marine carnivores (*Enhydra lutris*, *Lontra felina*, and *Ursus maritimus*). Bats do forage on land and have an impact, but we felt that they are too different to be reliably modelled along the rest – and therefore excluded them from out study. Further their ranges have not been documented to be affected by the expansion on the globe, and therefore wouldn’t change between current and present natural maps.

#### Densities

We estimated population density for all species. We used population densities from the PanTHERIA^53^ and body masses from PHYLACINE 1.2^34,35^ and underlying sources^54,55^. We build an allometric model of density as a function of body mass using a Bayesian approach based on species level phylogeny from PHYLACINE^34,35,56^, where estimates are weighted by closer phylogenetic relationships. Further explanation on the methods and overview of the imputed density results can be found in Appendix S1, with the estimates available in Table S2. In the further model we used the estimated densities for all species, to avoid biases by mixing known values and estimated values. To estimate the unbiased uncertainty we used 1000 of the sampled results for each species throughout the model.

Our consumption estimates are based on single population densities across a species range, which is fine for most species on average as there is no tendency for central abundance neither climatically of geographically^57^ – but estimates for individual species might be error prone. An example is the Capybara (*Hydrochoerus hydrochaeris*) a grazer which is highly connected to water, a resource not homogenously found across its large range^58,59^. Using a single density across its range limits our study’s geographic specific accuracy. We model density to 55/km^2^ – on par with an average (51/km^2^) previously found^60^, though densities between 1/km^2^ and 200/km^2^ have been reported^60^. Further, our models do not directly take species interactions into account, or density compensation by other species in a community if one goes extinct. Therefore, more species equates to more consumption in general. Some density compensation likely occurs in impoverished assemblages, while present-natural species assemblages probably more accurately reflect real densities and consumption.

#### Metabolic rates

We estimated field metabolic rates for all species. We first compiled a dataset of metabolic rates (MR) (Table S1). We gathered data on both basal metabolic rate (BMR) and field metabolic rate (FMR). FMR and BMR are closely related, and therefore known values of one could help pin down the value of the other estimate. FMR and BMR are were tightly relate to body mass even within each species, therefore we in the dataset often have several body mass/metabolic rate (MR) pairs for each species. We built an allometric model of MR as a function of body mass using a Bayesian approach based on the species-level phylogeny from PHYLACINE^34,35,56^, where estimates are weighted by closer phylogenetic relationships. Further explanation on the methods and overview of the imputed density results can be found in Appendix S2, with the estimates available in Table S3. In the further model we used the estimated densities based on the body masses from PHYLACINE for all species. To estimate the unbiased uncertainty we used 1000 of the sampled results for each species throughout the model.

#### Species energy needs and plant carbon consumption

For each species we calculated the species energy need (Equation 1), by multiplying FMR with density. By doing so we make the assumption that populations occur at stable densities across time, and that every animal eats at consumes the average needed energy for the species. This is assumption is wrong for young and suckling animals which require less energy and for pregnant and nursing animals which require more. On average these effects will at least partly cancel each other out.

Since not all species in our model only eat plants we corrected by the plant diet (Diet.Plant) percentage based on the PHYLACINE diet^34,35,61,62^. This will remove all strict non-plant eaters from impact, and reduce all not strict herbivores as well. Assuming that the plant diet percentage is equal to energy consumed is not necessarily correct, but plant diets percentages are calculated in a number of ways and a better solution is not available. And species affected by this counts only in the minority of the model anyway.

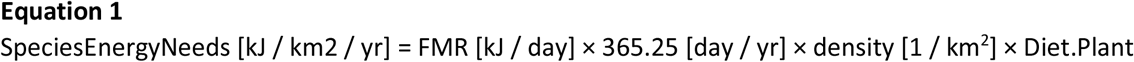

We were only interested in the animal’s impact on primary productivity and therefore weighted each species impact with its plant diet percentage (Equation 2). Several studies have shown that about 50% of the gross available energy is digested the rest is excreted again, and absorbed energy is ranges from 6.4-9.6 MJ/kgDM across wild monkeys, kangaroos and sheep^63–65^. We did not find enough species specific data for this factor, to differentiate between species. A study of diet selection in wild foraging sheep measured the metabolic energy (ME) available in the selected diet^65^. This study provided a measured of ME = 8.5 MJ / kg Dry-matter with an uncertainty (sd = 1.4). The uncertainty of the number was propagated through our models, by sampling this distribution (n = 1000). Further to translate kg Dry-matter to kg Carbon we need estimates for carbon content in vegetation. Vegetation contains about 45% (SD = 5.23) carbon (CC) [kgC / kgDM], which is does not vary much between plant ‘organ’ (fruit, stem, leaf, and root) or between life forms (e.g. herbs, broad-leaved trees, conifers, etc.), where the mean of any combination of life form and organ varied between 42% (herb-root) and 51% (conifer-stem)^66^. We used the mean vegetation CC for our model, and carried the uncertainty by sampling a normal distribution with the given parameters 1000 times.

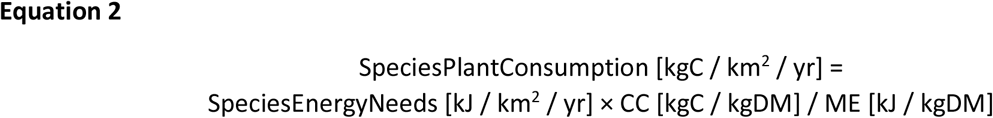

### Mapping the consumption

To compare estimate the human impact on natural consumption we used current range maps and present-natural range maps from PHYLACINE^6,34,35,52^. Present-natural range maps are counterfactual range maps with no mammal extinctions within the last 130 000 years. These maps are the baselines for all mapping the this paper, which are projected in a Berhman projection i.e. a ‘cylindrical equal’ area projection with standard parallel at 30° with a projected cell size of 96.5 km × 96.5 km, which is approximately 1° × 1° at 30° N and S.

For each grid cell we summed the species plant consumption for all species in that cell. By doing so we are making the assumption that each species occur at a uniform density across its full range. This is of course not the case, but basing it on any other distribution would introduce even more assumptions. Further, any comparisons between the current and the present natural will be minimally impacted by such an assumption. Still, the maps in PHYLACINE are very inclusive (i.e., if even a tiny part of a species range is in the cell, the whole cell is counted as range), and therefore we mitigate some of this by reducing species populations at range edges (defined using rook’s case). We do this based on the theoretical mean expectations if a grid is put on top of square ranges of random size. We focus on the number of rook’s case neighbours included in the range. Cells with no neighbors are weigted as 1/9^th^, endpoints (one neighbour) as 1/6^th^, bridges and corners (two neighbours) as 1/4^th^, and flat edges (three neighbours) as ½.

#### Global consumption of primary productivity

We compared our consumption estimates with primary productivity based on a mean dataset of corrected MODIS 17A3 NPP data from 2000-2015^67^. This was resampled to the same projection as the range maps. Species were scored as presence/absence within each cell (except for ½ at the range boundaries), which is generally justifiable for relatively homogeneous areas but it is increasingly problematic for more heterogeneous areas. We therefore removed the cells with most variable NPP from the analysis. Specifically we calculated the standard deviation of log_10_(NPP(g Carbon/m^2^/yr)+1) within each cell and excluded the upper 5% quantile of cells. Further we believe our estimates of mammal densities to be highly inaccurate in low production areas and therefore removed them, which has also been done in similar studies^21^. We defined low production as cells with NPP < 200 g Carbon/m^2^/yr, which is equivalent to most (¾-quantile) of the WWF biome Deserts and Xeric Shrublands a balance between not removing too much or too little.

Our estimates do not take the domestic mammals into account, or current modern land use. Therefore our numbers are solely estimated densities that are not farmed or domestically grazed – or in other ways influenced by human use. I.e. our current maps estimates densities in natural areas where population sizes are at their natural equilibrium for the species still present in the areas. Further we assume that all species occur across each full cell (except range edges where we assume 50%), and that they are not affected by poorer habitat. Finally we note that we are only mapping species within their native ranges and e.g. does not include the wild dromedaries in Australia^68^ since no systematic range estimate is available mapping the introduced ranges of all mammals.

To summarise our findings across the wildest remaining places on earth, we downloaded the ‘Last of the Wild’^37,50^ dataset (Downloaded from http://sedac.ciesin.columbia.edu/data/set/wildareas-v2-last-of-the-wild-geographic). This map was resampled and projected to match the maps from PHYLACINE 1.2, and extracted to ecoregions and biomes c.f. WWF^51^. The map includes all the areas with a human footprint of maximum 10 (out of 100)^50^.

## Supporting information

Table S2b Imputed density

Table S3b Imputed metabolic rates

## Acknowledgements

We thank the VILLUM Fonden and the Carlsberg Foundation for economic support, through a VILLUM Investigator grant for the project “Biodiversity Dynamics in a Changing World” (grant 16549, to J.-C-S.) and a Semper Ardens grant for the project MegaPast2Future (grant CF16-0005, to J.-C-S.). We furthermore thank the European Research Council for economic support (ERC-2012-StG-310886-HISTFUNC, to J.-C-S.), S.F. was supported by the Swedish research council (2017-03862).

We assembled data mostly from larger databases and it was therefore not possible to cite all the original underlying studies, but we are grateful for all the preceding efforts in primary data collection and in data integration and harmonization, which enabled us to carry out this study.

## Supplementary materials overview

### Appendices

Appendix S1: Imputation of density

Appendix S2: Imputation of metabolic rate

**Table S1a:** Metabolic rate data. Column explanations.

| Column name | Column explanation |
| --- | --- |
| Binomial.1.2 | Phylacine Taxonomy |
| Order.1.2 | Phylacine Taxonomy |
| Family.1.2 | Phylacine Taxonomy |
| Binomial.Source | Binomial used in source |
| BM | Mass for measured individual in gram |
| MR | Metabolic rate kJ/day |
| log10BM | Log10 Mass for measured individual in gram |
| log10MR | Log10 Metabolic rate kJ/day |
| MR.type | Basal or field metabolic rate (BMR/FMR) |
| Source | Reference to source |
| Comment | Comment for single species sources |

**Table S1b:** Metabolic rate data.

**Table S2a:**
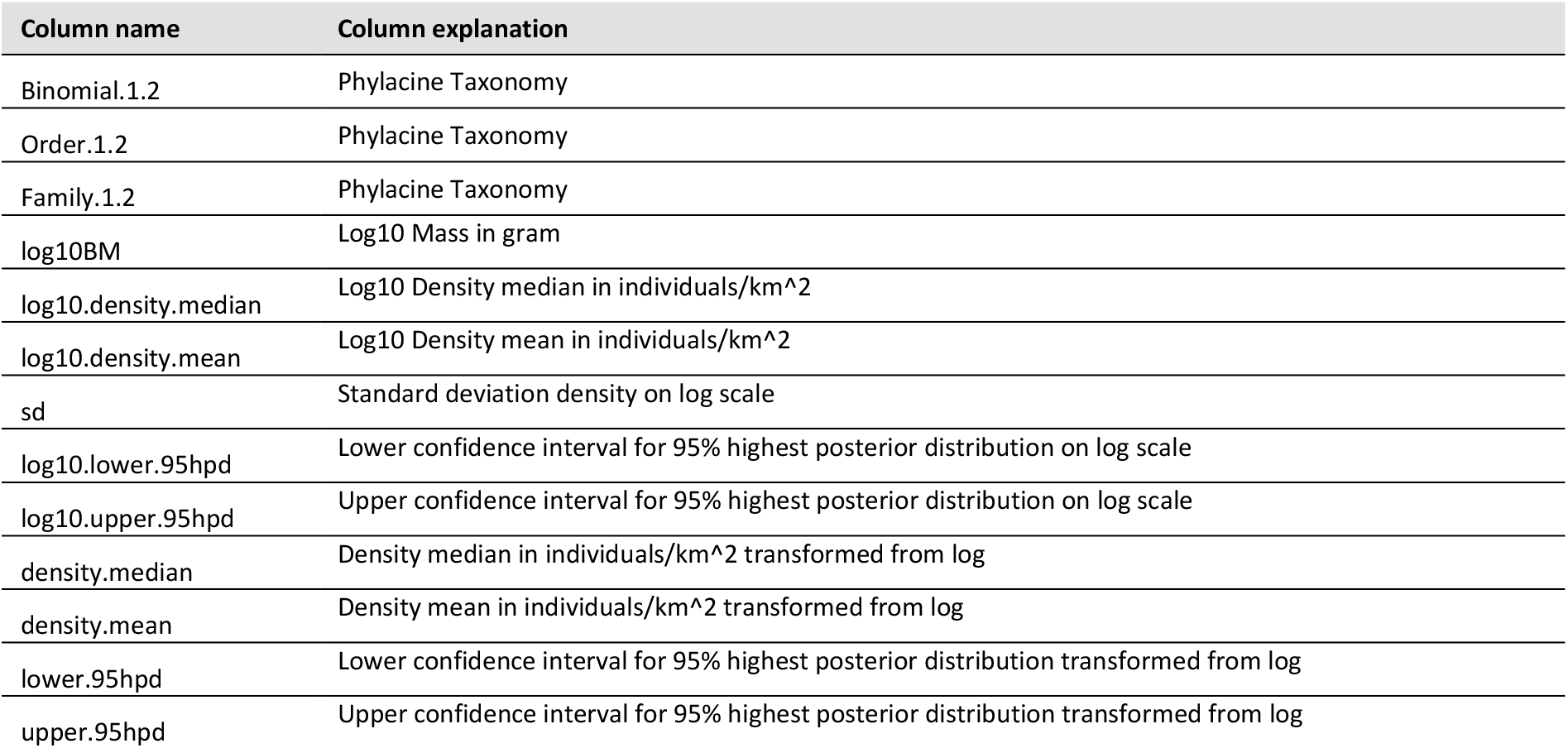
Imputed density. Column explanations.

**Table S2b:** Imputed density.

**Table S3a:**
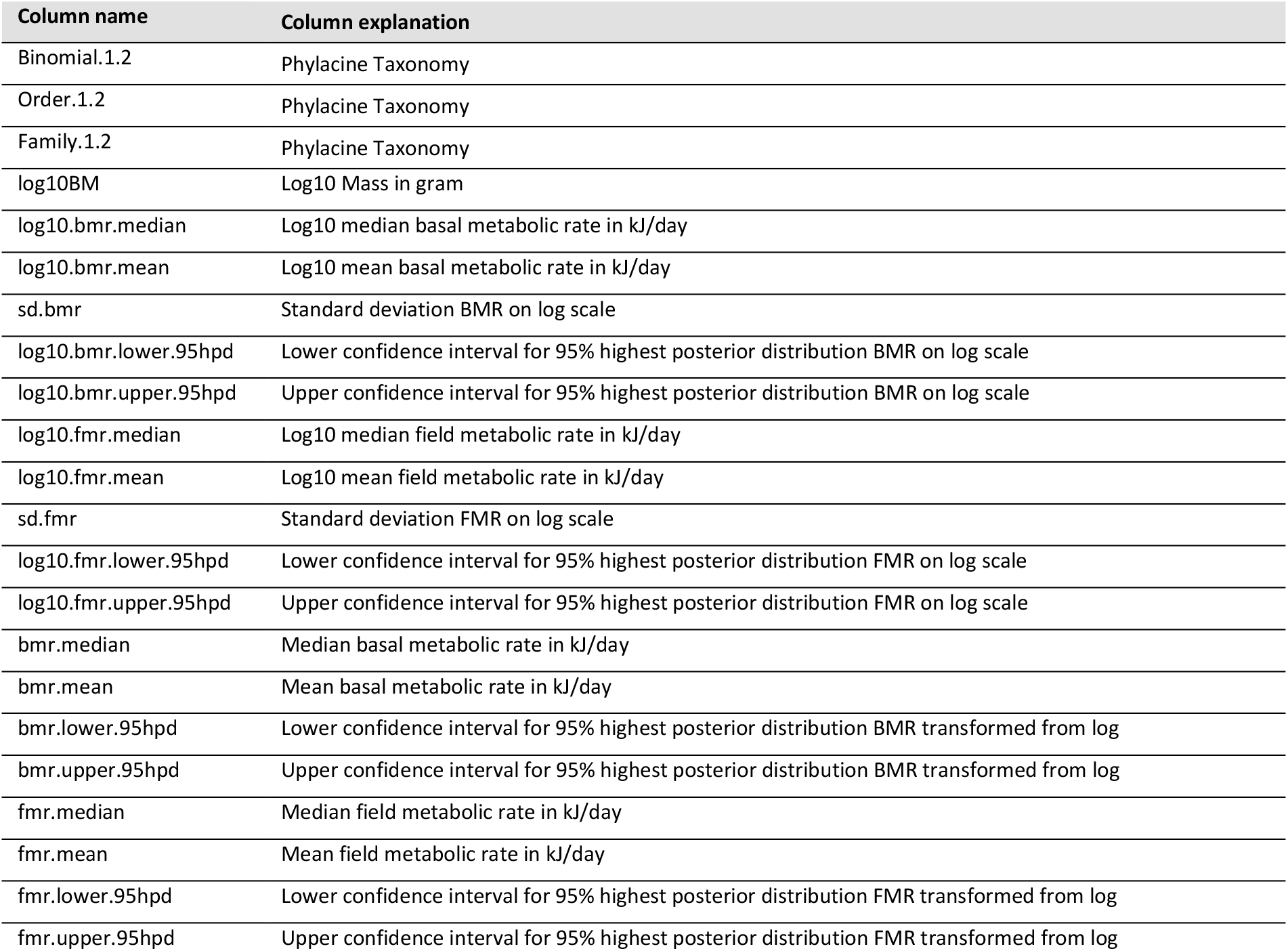
Imputed metabolic rate. All estimates are for the average species body mass from PHYLACINE. Column explanations.

**Table S3b:** Imputed metabolic rate. All estimates are for the average species body mass from PHYLACINE.

**Fig S1:**
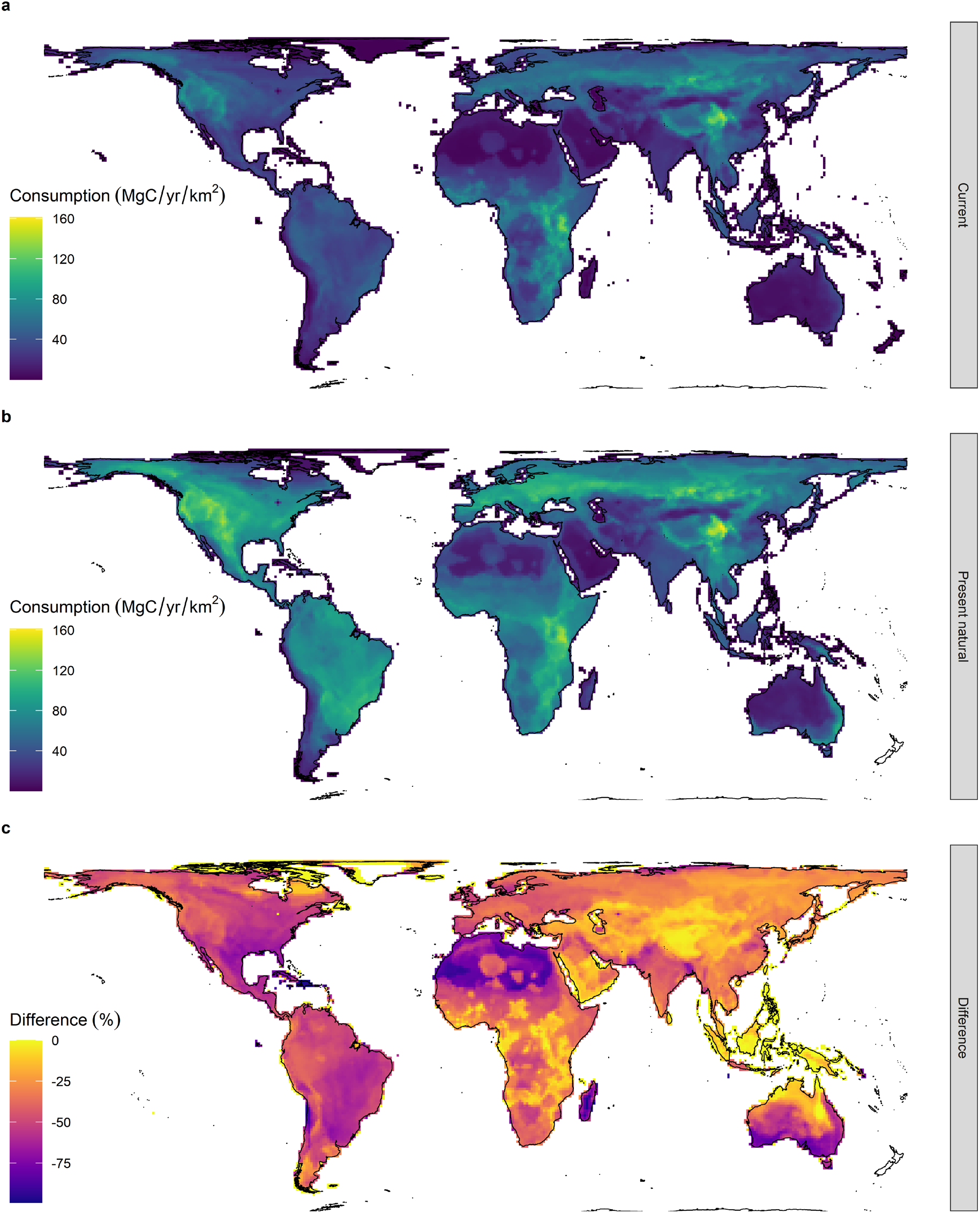
Mammal consumption of plant carbon globally. Same as Fig 1, but no areas removed. a) Average terrestrial mammal consumption by current mammals (current ranges of extant species). b) shows present natural consumption (potential present day ranges with no human presence). c) shows the percentage lower consumption of current ranges compared to present natural ranges.

**Fig S2:**
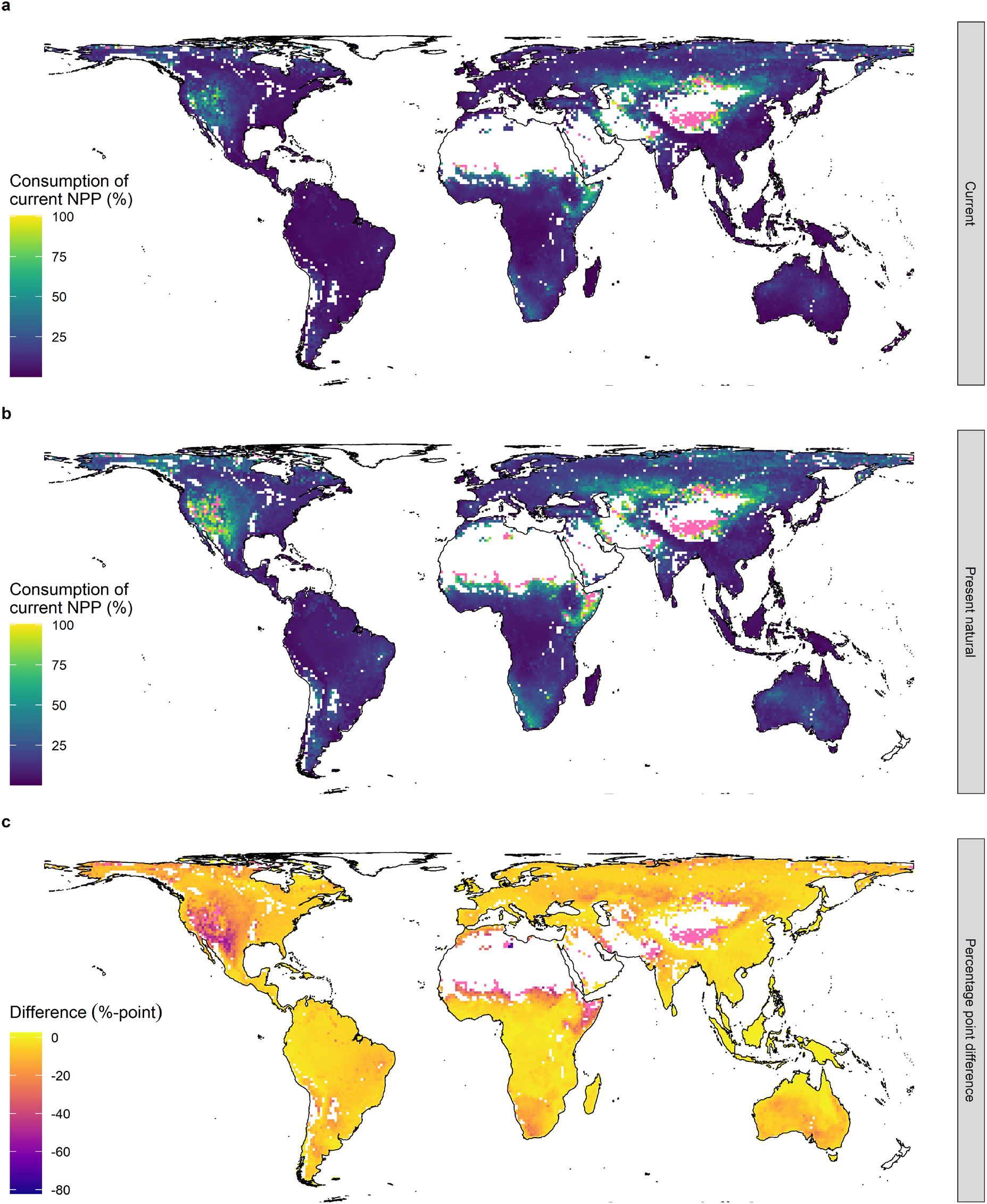
Estimated mammal consumption of net primary productivity (NPP) globally. Same as Fig 2 but including the low NPP areas where our model likely causes overestimations of densities. a) Consumption by current mammals (current ranges of extant species). b) Present natural consumption (potential present day ranges with no human presence). c) Percentage-point consumption difference between present natural and current consumption. Blank pixels are either unknown or high variability in NPP and pixels in hot pink are where consumptions exceeds 100%.

**Fig S3:**
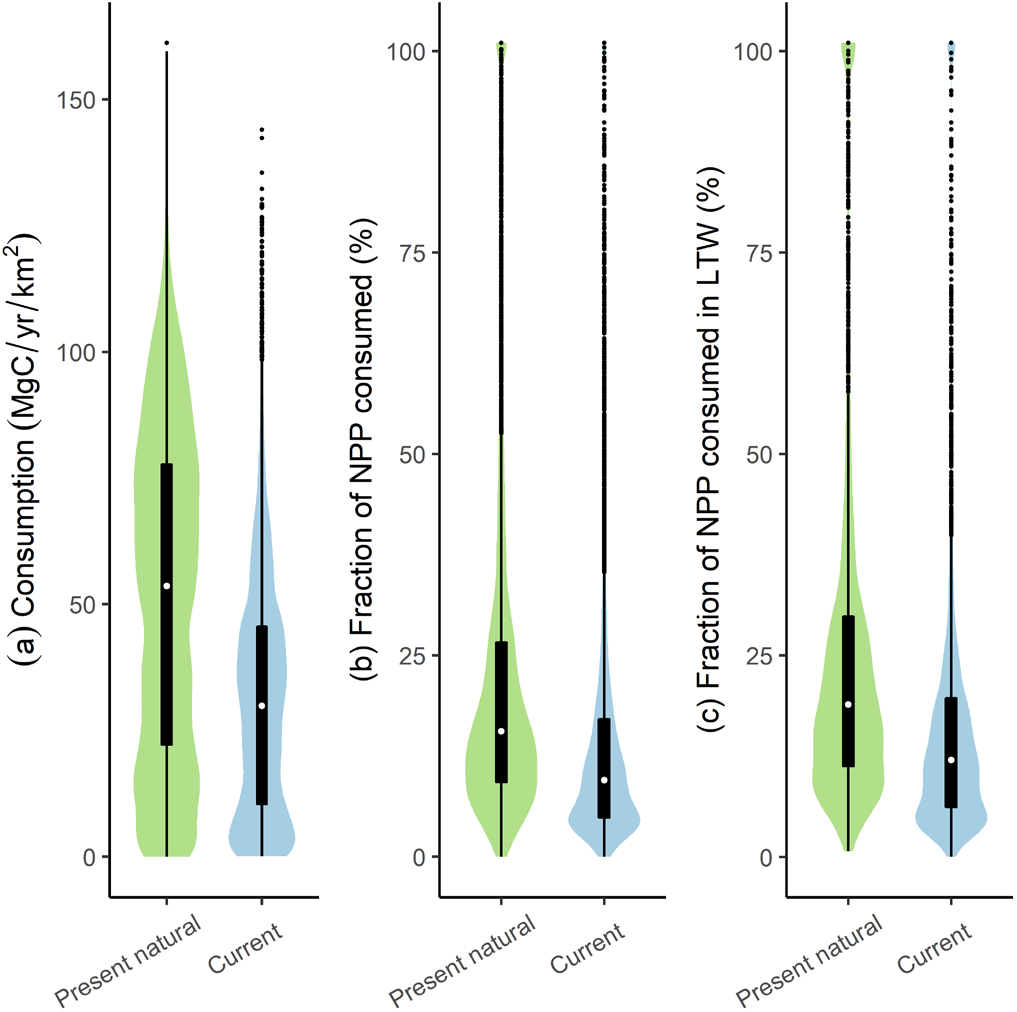
Global consumption summary. Same as Fig 3, but low NPP areas not removed. Boxplot with underlying violins with kernel density scaled to width. **a)** Total consumption of carbon. **b)** Fraction of net primary productivity (NPP) consumed. **c)** Fraction of net primary productivity (NPP) consumed in the areas designated as ‘last of the wild’^37,50^. Values above 100% are truncated to 101%.

**Fig S4:**
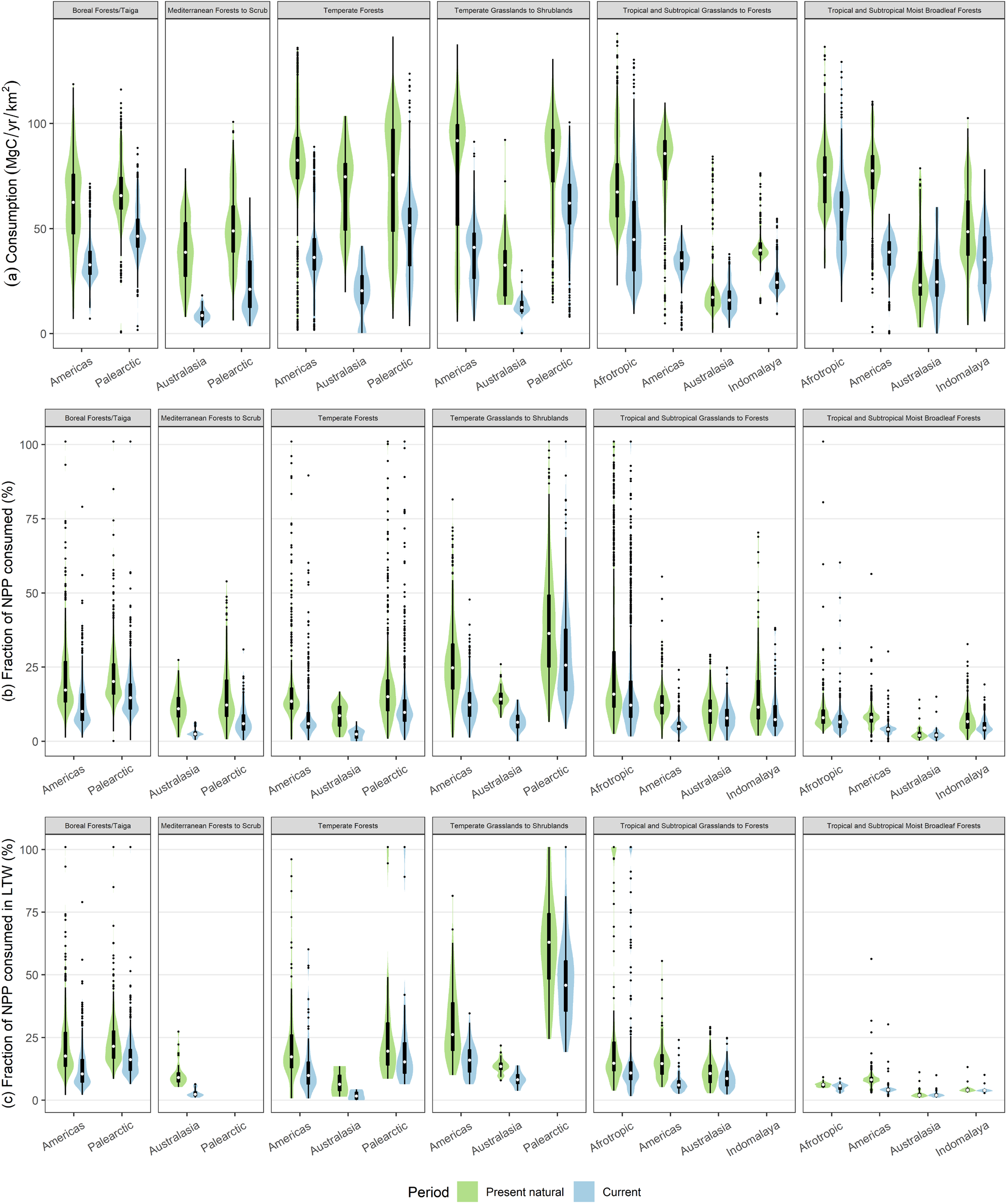
Consumption summary for large ecological units based on realms and biomes^51^. Same as Fig 4, but low NPP areas not removed. Boxplot with underlying violins with kernel density scaled to width. Neotropics and Nearctic are merged to the Americas and Madagascar is excluded from the Afrotropics for biome modifications see Fig S5. **a)** Total consumption of carbon. **b)** Fraction of net primary productivity (NPP) consumed. **c)** Fraction of net primary productivity (NPP) consumed in the areas designated as ‘last of the wild’^37,50^. Values above 100% are truncated to 101%.

**Fig S5:**
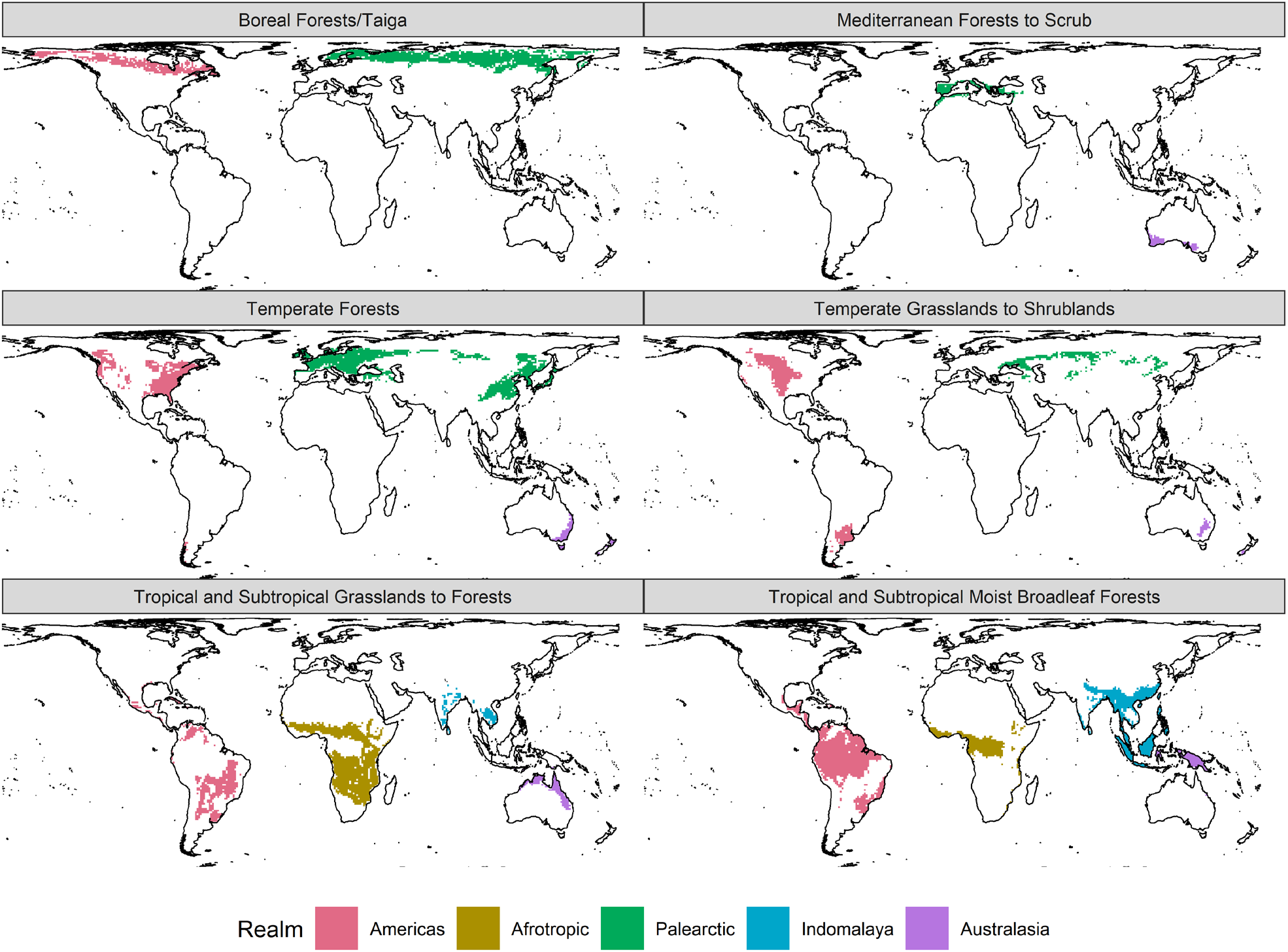
Global ecological units based on realms and biomes^51^. Neotropics and Nearctic are merged in the Americas and Madagascar is excluded from the Afrotropics. “*Temperate Broadleaf and Mixed Forests”* and “*Temperate Coniferous Forests”* are merged to “*Temperate Forests”*. “*Tropical and Subtropical Dry Broadleaf Forests”* and “*Tropical and subtropical grasslands, savannas, and shrublands”* are merged to “*Tropical and Subtropical Grasslands to Forests”*. We changed the name of “*Temperate Grasslands, Savannas, and Shrublands”* to “*Temperate Grasslands to Shrublands”*, and changed “*Mediterranean Forests, Woodlands, and Scrub”* to “*Mediterranean Forests to Scrub”* for plotting purposes.

**Fig S6:**
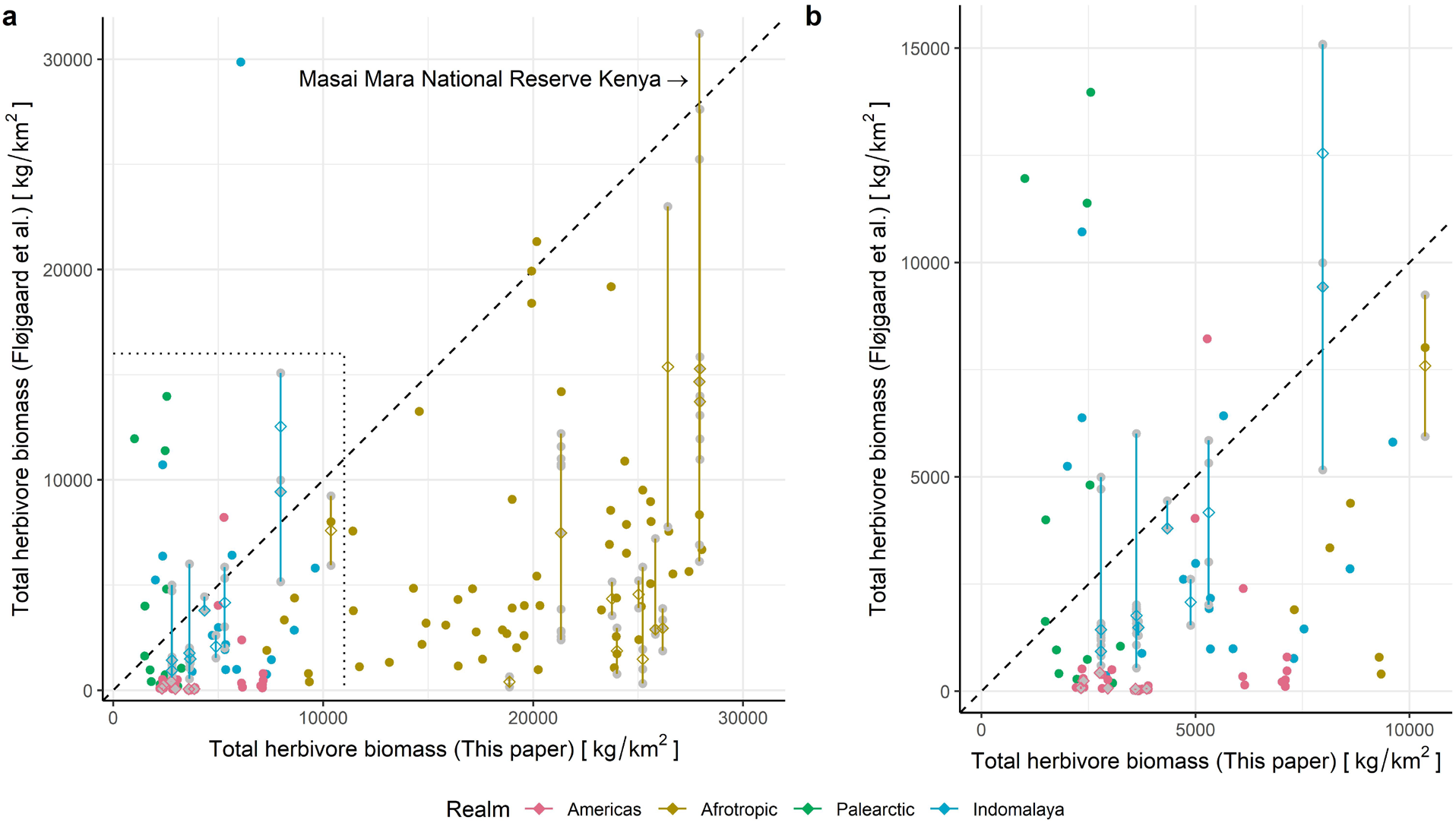
Total current biomass in protected areas. Data of herbivore biomass estimates based on population counts from Fløjgaard et al.^24^ mapped against this study’s median results in the same areas (for their current fauna). (a) Is all global data and (b) is a zoomed region showing more detailed info outside the Afrotropics. Areas that have been estimated across several years are shown as line-ranges with median diamonds and grey points for yearly estimates. In areas where there are multiple estimates from Fløjgaard et al. we often see that our estimate is within the observed range. The Maasai Mara National Reserve in Kenya (line) have been highlighted as an example of an area with close to intact megafauna.

## References

1. Sandom, C. J., Ejrnaes, R., Hansen, M. D. D. D. & Svenning, J.-C. J. C. High herbivore density associated with vegetation diversity in interglacial ecosystems. Proc. Natl. Acad. Sci. 111, 4162–4167 (2014).

2. Daskin, J. H., Stalmans, M. & Pringle, R. M. Ecological legacies of civil war: 35-year increase in savanna tree cover following wholesale large-mammal declines. J. Ecol. 104, 79–89 (2016).

3. Bakker, E. S. et al. Combining paleo-data and modern exclosure experiments to assess the impact of megafauna extinctions on woody vegetation. Proc. Natl. Acad. Sci. 113, 847–855 (2016).

4. Cebrian, J. & Lartigue, J. Patterns of herbivory and decomposition in aquatic and terrestrial ecosystems. Ecol. Monogr. 74, 237–259 (2004).

5. Ripple, W. J. et al. Collapse of the world’s largest herbivores. Sci. Adv. 1, e1400103–e1400103 (2015).

6. Faurby, S. & Svenning, J.-C. Historic and prehistoric human-driven extinctions have reshaped global mammal diversity patterns. Divers. Distrib. 21, 1155–1166 (2015).

7. Sandom, C., Faurby, S., Sandel, B. & Svenning, J.-C. Global late Quaternary megafauna extinctions linked to humans, not climate change. Proc. R. Soc. B Biol. Sci. 281, 20133254 (2014).

8. Andermann, T., Faurby, S., Turvey, S. T., Antonelli, A. & Silvestro, D. The past and future human impact on mammalian diversity. Sci. Adv. 6, eabb2313 (2020).

9. Williams, R., Bradstock, R. & Cary, G. Interactions between climate change, fire regimes and biodiversity in Australia: a preliminary assessment. (2009).

10. Asner, G. P. & Levick, S. R. Landscape-scale effects of herbivores on treefall in African savannas. Ecol. Lett. 15, 1211–1217 (2012).

11. Munn, A. J., Dunne, C., Müller, D. W. H. & Clauss, M. Energy In-Equivalence in Australian Marsupials: Evidence for Disruption of the Continent’s Mammal Assemblage, or Are Rules Meant to Be Broken? PLoS One 8, 1–5 (2013).

12. Pedersen, R. Ø., Faurby, S. & Svenning, J.-C. Shallow size-density relations within mammal clades suggest greater intra-guild ecological impact of large-bodied species. J. Anim. Ecol. 86, 1205–1213 (2017).

13. Doughty, C. E., Faurby, S. & Svenning, J.-C. The impact of the megafauna extinctions on savanna woody cover in South America. Ecography (Cop.). 39, 213–222 (2016).

14. Rule, S. et al. The Aftermath of Megafaunal Extinction: Ecosystem Transformation in Pleistocene Australia. Science (80-.). 335, 1483–1486 (2012).

15. Gill, J. L., Williams, J. W., Jackson, S. T., Lininger, K. B. & Robinson, G. S. Pleistocene Megafaunal Collapse, Novel Plant Communities, and Enhanced Fire Regimes in North America. Science (80-.). 326, 1100–1103 (2009).

16. Guyton, J. A. et al. Trophic rewilding revives biotic resistance to shrub invasion. Nat. Ecol. Evol. 4, 712–724 (2020).

17. Beschta, R. L., Ripple, W. J., Kauffman, J. B. & Painter, L. E. Bison limit ecosystem recovery in northern Yellowstone. Food Webs 23, e00142 (2020).

18. Churski, M., Bubnicki, J. W., Jędrzejewska, B., Kuijper, D. P. J. & Cromsigt, J. P. G. M. Brown world forests: increased ungulate browsing keeps temperate trees in recruitment bottlenecks in resource hotspots. New Phytol. 214, 158–168 (2017).

19. Davies, A. B., Gaylard, A. & Asner, G. P. Megafaunal effects on vegetation structure throughout a densely wooded African landscape. Ecol. Appl. 28, 398–408 (2018).

20. Donlan, J. et al. Pleistocene rewilding: an optimistic agenda for twenty-first century conservation. Am. Nat. 168, 660–81 (2006).

21. Doughty, C. E., Faurby, S., Wolf, A., Malhi, Y. & Svenning, J.-C. C. Changing NPP consumption patterns in the Holocene: From megafauna-‘liberated’ NPP to ‘ecological bankruptcy’. Anthr. Rev. 3, 174–187 (2016).

22. Barnosky, A. D. In the Light of Evolution. In the Light of Evolution 2, (National Academies Press, 2008).

23. Bar-On, Y. M., Phillips, R. & Milo, R. The biomass distribution on Earth. Proc. Natl. Acad. Sci. U. S. A. 115, 6506–6511 (2018).

24. Fløjgaard, C., Pedersen, P. B., Sandom, C., Svenning, J.-C. & Ejrnæs, R. Exploring a natural baseline for large herbivore biomass. 1–18 (2020). doi:10.1101/2020.02.27.968461

25. Robson, A. S. et al. Savanna elephant numbers are only a quarter of their expected values. PLoS One 12, e0175942 (2017).

26. Ogutu, J. O. et al. Extreme Wildlife Declines and Concurrent Increase in Livestock Numbers in Kenya: What Are the Causes? PLoS One 11, e0163249 (2016).

27. Ogutu, J. O., Owen-Smith, N., Piepho, H.-P. & Said, M. Y. Continuing wildlife population declines and range contraction in the Mara region of Kenya during 1977-2009. J. Zool. 285, 99–109 (2011).

28. McNaughton, S. J. Ecology of a Grazing Ecosystem: The Serengeti. Ecol. Monogr. 55, 259–294 (1985).

29. Lundgren, E. J. et al. Introduced herbivores restore Late Pleistocene ecological functions. Proc. Natl. Acad. Sci. 117, 7871–7878 (2020).

30. Poveda, K., Steffan-Dewenter, I., Scheu, S. & Tscharntke, T. Effects of below- and above-ground herbivores on plant growth, flower visitation and seed set. Oecologia 135, 601–605 (2003).

31. Belovsky, G. E. & Slade, J. B. Insect herbivory accelerates nutrient cycling and increases plant production. Proc. Natl. Acad. Sci. 97, 14412–14417 (2000).

32. Zimov, S. A., Zimov, N. S., Tikhonov, A. N. & Chapin, I. S. Mammoth steppe: A high-productivity phenomenon. Quat. Sci. Rev. 57, 26–45 (2012).

33. Semmartin, M., Oesterheld, M., Semmartin, M. & Oesterheld, M. Effect of Grazing Pattern on Primary Productivity. Oikos 75, 431 (1996).

34. Faurby, S. et al. PHYLACINE 1.2.1: An update to the Phylogenetic Atlas of Mammal Macroecology. (2020). doi:10.5281/zenodo.3690867

35. Faurby, S. et al. PHYLACINE 1.2: The Phylogenetic Atlas of Mammal Macroecology. Ecology 99, 2626–2626 (2018).

36. Pedersen, R. Ø. & Davis, M. PHYLACINE 1.2 Vignette. (2018). Available at: https://github.com/MegaPast2Future/PHYLACINE_1.2#vignette. (Accessed: 11th October 2020)

37. WCS, W. C. S.- & University, C. for I. E. S. I. N.-C.-C. Last of the Wild Project, Version 2, 2005 (LWP-2): Last of the Wild Dataset (Geographic). (2005). doi:10.7927/H4348H83

38. Belote, R. T. et al. Mammal species composition reveals new insights into Earth’s remaining wilderness. Front. Ecol. Environ. 1–8 (2020). doi:10.1002/fee.2192

39. Sandom, C. J. et al. Trophic rewilding presents regionally specific opportunities for mitigating climate change. Philos. Trans. R. Soc. B Biol. Sci. 375, 20190125 (2020).

40. Gill, J. L. Ecological impacts of the late Quaternary megaherbivore extinctions. New Phytol. 201, 1163–1169 (2014).

41. Berti, E. & Svenning, J. Megafauna extinctions have reduced biotic connectivity worldwide. Glob. Ecol. Biogeogr. geb.13182 (2020). doi:10.1111/geb.13182

42. Peco, B., Sánchez, A. M. & Azcárate, F. M. Abandonment in grazing systems: Consequences for vegetation and soil. Agric. Ecosyst. Environ. 113, 284–294 (2006).

43. Damschen, E. I. et al. Ongoing accumulation of plant diversity through habitat connectivity in an 18-year experiment. Science (80-.). 365, 1478–1480 (2019).

44. Johnson, C. N. Ecological consequences of late quaternary extinctions of megafauna. Proc. R. Soc. B Biol. Sci. 276, 2509–2519 (2009).

45. Doughty, C. E., Wolf, A. & Field, C. B. Biophysical feedbacks between the Pleistocene megafauna extinction and climate: The first human-induced global warming? Geophys. Res. Lett. 37, n/a–n/a (2010).

46. Guldemond, R. A. R., Purdon, A. & van Aarde, R. J. A systematic review of elephant impact across Africa. PLoS One 12, e0178935 (2017).

47. Davis, M. What North America’s skeleton crew of megafauna tells us about community disassembly. Proc. R. Soc. B Biol. Sci. 284, 20162116 (2017).

48. Cooke, R. S. C., Eigenbrod, F. & Bates, A. E. Projected losses of global mammal and bird ecological strategies. Nat. Commun. 10, 2279 (2019).

49. Svenning, J.-C. et al. Science for a wilder Anthropocene: Synthesis and future directions for trophic rewilding research. Proc. Natl. Acad. Sci. 113, 898–906 (2016).

50. Sanderson, E. W. et al. The Human Footprint and the Last of the Wild. Bioscience 52, 891 (2002).

51. Olson, D. M. et al. Terrestrial Ecoregions of the World: A New Map of Life on Earth. Bioscience 51, 933 (2001).

52. IUCN. IUCN Red List of Threatened Species. Version 2015.4 www.iucnredlist.org (2015). Available at: www.iucnredlist.org. (Accessed: 1st January 2015)

53. Jones, K. E. et al. PanTHERIA: a species-level database of life history, ecology, and geography of extant and recently extinct mammals. Ecology 90, 2648–2648 (2009).

54. Smith, F. A. et al. Body mass of late quaternary mammals. Ecology 84, 3403–3403 (2003).

55. Faurby, S. & Svenning, J.-C. Resurrection of the Island Rule: Human-Driven Extinctions Have Obscured a Basic Evolutionary Pattern. Am. Nat. 187, 812–820 (2016).

56. Faurby, S. & Svenning, J.-C. A species-level phylogeny of all extant and late Quaternary extinct mammals using a novel heuristic-hierarchical Bayesian approach. Mol. Phylogenet. Evol. 84, 14–26 (2015).

57. Morgan, D., Sanz, C., Onononga, J. R. & Strindberg, S. Factors Influencing the Survival of Sympatric Gorilla (Gorilla gorilla gorilla) and Chimpanzee (Pan troglodytes troglodytes) Nests. Int. J. Primatol. 37, 718–737 (2016).

58. IUCN. The IUCN Red List of Threatened Species. Version 2016-3 (2016). Available at: http://www.iucnredlist.org.

59. Wilson, D. E., Lacher, T. E. & Mittermeier, R. A. Handbook of the mammals of the World. Vol. 6 Lagomorphs and Rodents I. Lynx Edicions (2016).

60. Verdade, L. M. & Ferraz, K. M. P. M. B. Capybaras in an anthropogenic habitat in Southeastern Brazil. Brazilian J. Biol. 66, 371–378 (2006).

61. Wilman, H. et al. EltonTraits 1.0: Species-level foraging attributes of the world’s birds and mammals. Ecology 95, 2027–2027 (2014).

62. Kissling, W. D. et al. Establishing macroecological trait datasets: digitalization, extrapolation, and validation of diet preferences in terrestrial mammals worldwide. Ecol. Evol. 4, 2913–30 (2014).

63. Nagy, K. A., Milton, K., Nagy, K. A. & Milton, K. Energy Metabolism and Food Consumption by Wild Howler Monkeys. 60, 475–480 (2016).

64. Munn, A. J., Dawson, T. J., McLeod, S. R., Dennis, T. & Maloney, S. K. Energy, water and space use by free-living red kangaroos Macropus rufus and domestic sheep Ovis aries in an Australian rangeland. J. Comp. Physiol. B Biochem. Syst. Environ. Physiol. 183, 843–858 (2013).

65. Degen, A. A., Benjamin, R. W., Abdraimov, S. A. & Sarbasov, T. I. Browse selection by Karakul sheep in relation to plant composition and estimated metabolizable energy content. J. Agric. Sci. 139, 353–358 (2002).

66. Ma, S. et al. Variations and determinants of carbon content in plants: A global synthesis. Biogeosciences 15, 693–702 (2018).

67. Zhao, M., Heinsch, F. A., Nemani, R. R. & Running, S. W. Improvements of the MODIS terrestrial gross and net primary production global data set. Remote Sens. Environ. 95, 164–176 (2005).

68. Lundgren, E. J., Ramp, D., Ripple, W. J. & Wallach, A. D. Introduced megafauna are rewilding the Anthropocene. Ecography (Cop.). 857–866 (2017). doi:10.1111/ecog.03430

