## Appendix S1 for "Extinctions have strongly reduced the mammalian consumption of primary productivity"

### Population densities

In the following we provide our method for imputing mammal population densities, and provide a full summarised dataset of the results.

#### Data

For species densities we used all available data from PanTHERIA (Jones et al., 2009), and for phylogeny we used a posterior distribution of 1000 equally likely fully resolved phylogenies for all mammals provided with the PHYLACINE (v. 1.2.1) database (Faurby et al., 2018, 2020; Faurby & Svenning, 2015). Body mass was also acquired from PHYLACINE (v. 1.2.1) and underlying sources (Faurby et al., 2018, 2020; Faurby & Svenning, 2016). The dataset was filtered to species classified as non-marine, non-flying, terrestrial species (i.e. excluding bats, sea cows, whales, pinnipeds and three mostly marine carnivores *Enhydra lutris*, *Lontra felina*, and *Ursus maritimus*. The final dataset of mass and population density consisted of 929 species spanning 21 orders and 97 families, well distributed across the phylogeny and mass range (Fig 2).

#### Taxonomy

Species names were aligned to PHYLACINE, using the synonymy table provided in PHYLACINE, 2 species which did not resolve were corrected manually *Cercopithecus pogonias* and *Cebus olivaceus* were recognized as *Cercopithecus denti* and *Cebus brunneus* respectively. Data for 3 species (*Damaliscus lunatus*, *Felis silvestris*, and *Alcelaphus buselaphus*) cf. PHYLACINE had duplicated records in PanTHERIA and only records with the nominal species were kept and the other species records were disregarded (*Damaliscus korrigum*, *Felis catus*, *Alcelaphus caama*, and *Alcelaphus lichtensteinii*).

#### Imputation

We used the 'MCMCglmm' package v. 2.29 (J. D. Hadfield, 2010) in the R language v. 3.6.2 (R Core Team, 2017) for imputation, where we used relatively uninformative priors as recommended (J. Hadfield, 2012), G(V = 1, nu = 0.002) and R(V = 1, nu = 0.002). The model was specified as $\log_{10}$ *Density* as a function of $\log_{10}$ *Body mass* with the phylogeny as random effect. We found a burnin of 3000 with a thinning factor of 100 to be sufficient, which ran for 36 300 iterations on 3 chains giving 999 samples per species for one test tree. Effective sample size for intercept was 298 and slope was 333 per chain, and 128 for the G-structure on phylogeny and 239 for the R-structure (Fig 4 and 5), Gelman diagnostics value was found < 1.02 for all variables. For the full dataset of trees only 3 samples per chain was necessary to give an overall impression of the value distribution across the 1000 trees. Across a posterior distribution of 1000 equally likely phylogenetic trees from PHYLACINE, giving 9000 samples per species.

The following packages was used: 'doSNOW' v. 1.0.18 (Corporation & Weston, 2017) for parallel processing, 'ape' v. 5.3 (Paradis, Claude, & Strimmer, 2004) for handling phylogenies, and 'tidyverse' v. 1.3.0 (Wickham, 2017) for general data wrangling and plotting.

#### Results

The final results were summarised in two ways. The final dataset with 9000 samples for each species density was used throughout our calculations, and summarised everything with mean, median, and Highest Posterior Distribution (HPD)-95% credibility interval in Table S2b. We here provide only the estimated values with their uncertainties and exclude the values from PanTHERIA for a uniform dataset (Fig 1-2). Estimated densities are distributed evenly around the empirical data and values and are relatively unbiased across the dataset, though with a tendency for overestimates of small densities and underestimates of large densities (Fig 3). This is not biased across body masses (Fig 4).

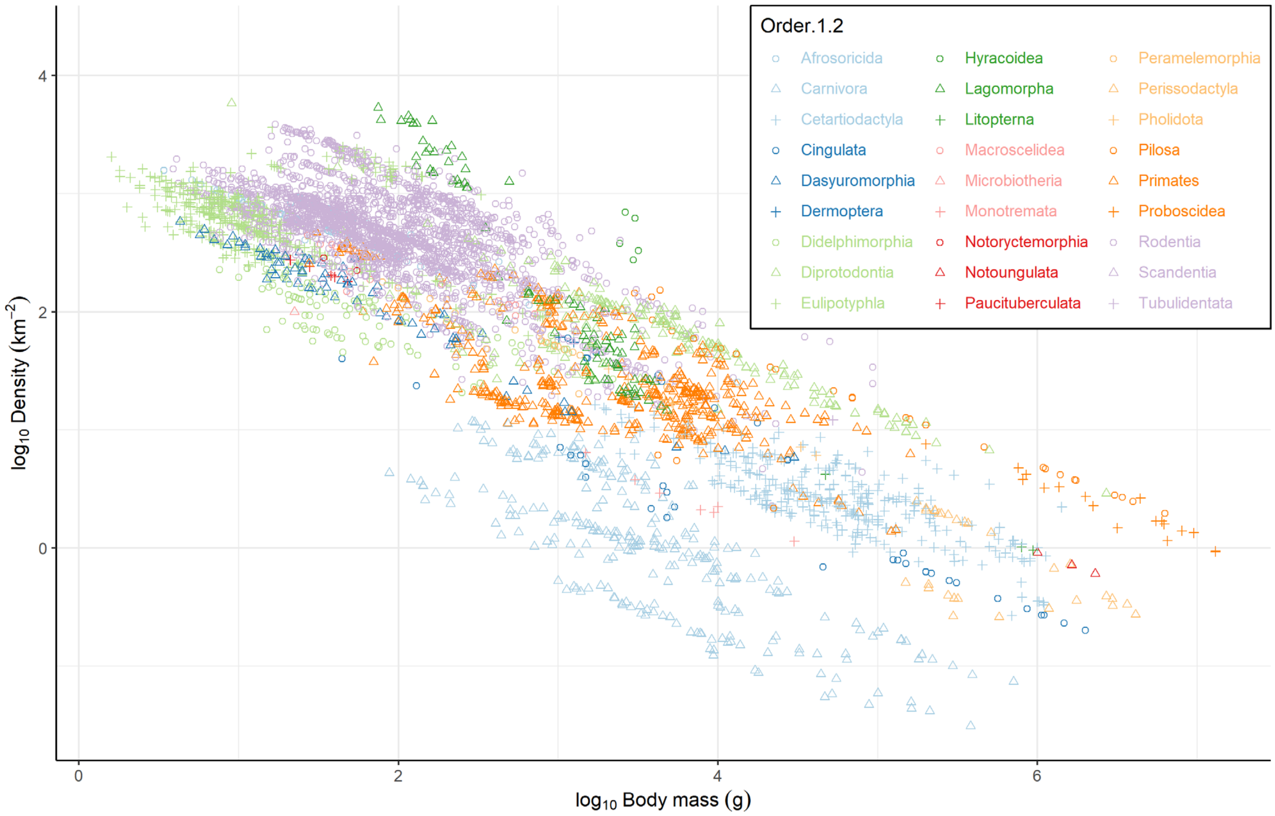


**Fig 1:** Median density estimates for all terrestrial mammals plotted along body size on a log10-log10 scale. The 29 orders of mammals are coded by a unique combination of symbol and colour to distinguish them as much as possible.

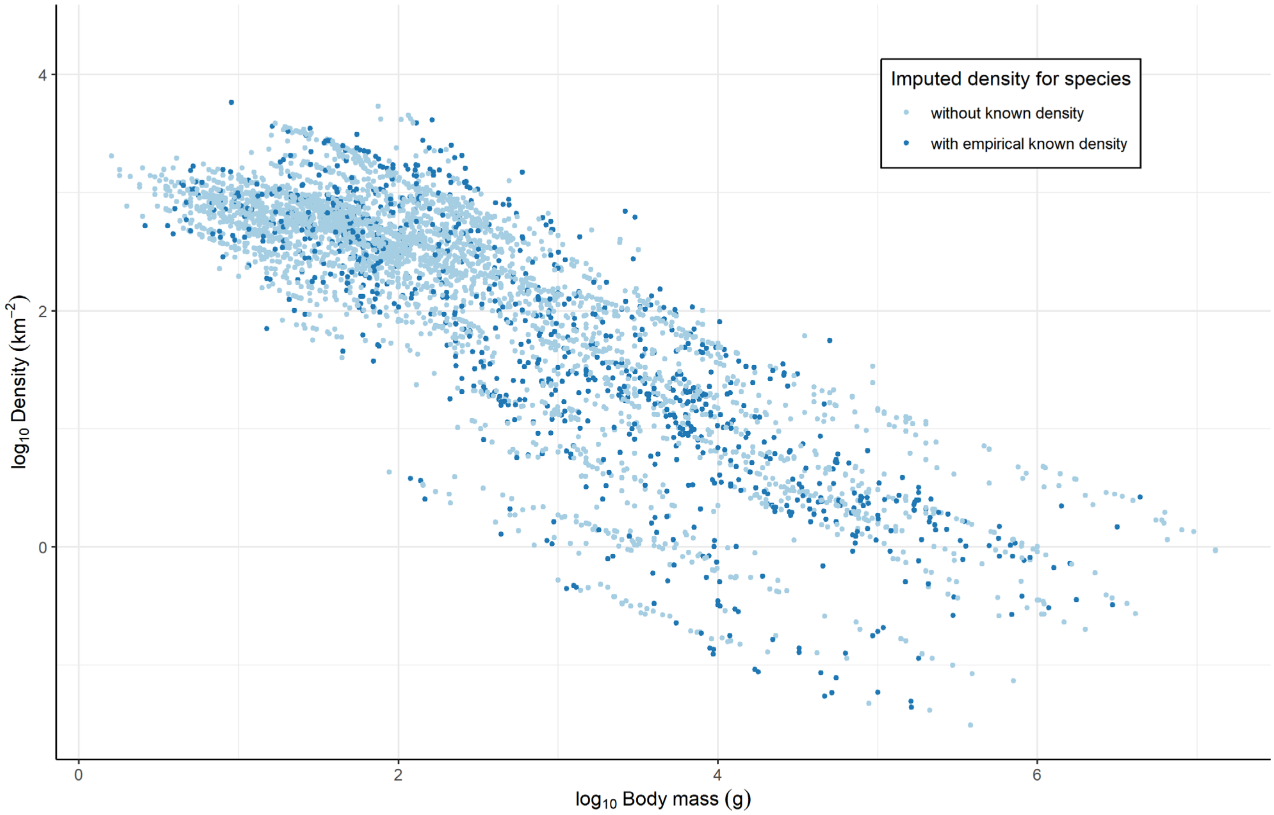


**Fig 2:** Mean imputed densities for all terrestrial mammals. All values are imputed, orange points indicate species for which we have empirical density estimates in our model.

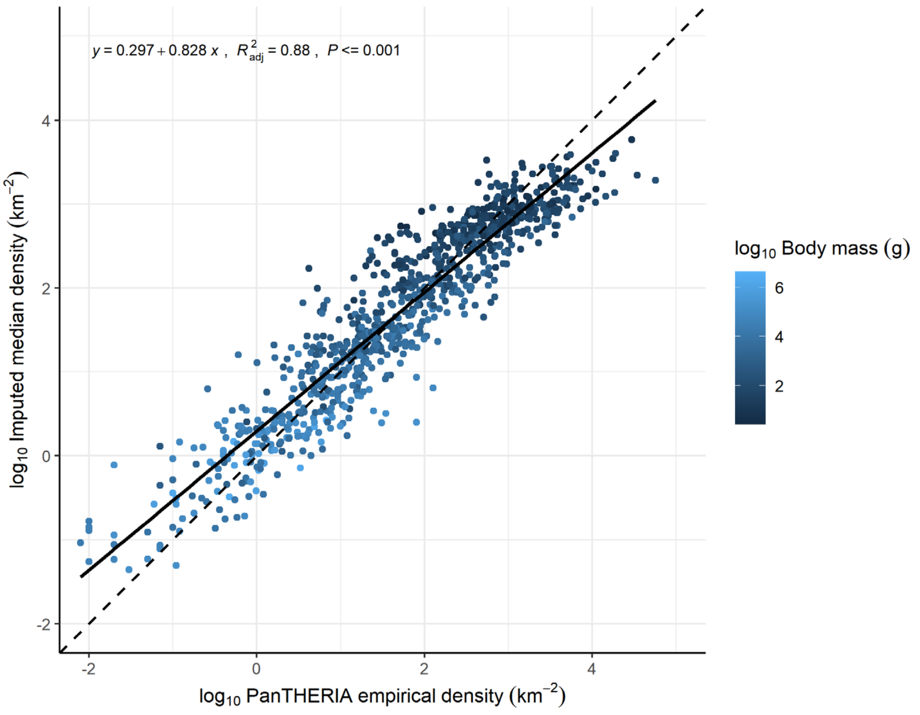


**Fig 3:** Overview of the imputed values compared to the empirical estimates from PanTHERIA. Color brightness shows body mass, the dashed line shows the 1:1 line, and the solid line indicates a linear fit. The plot shows a spread of values of the imputed densities about a factor 10 around the empirical values. On average the model seems have slightly over-estimated small population densities and underestimated large population densities. This effect is not driven by body mass bias (Fig 4).

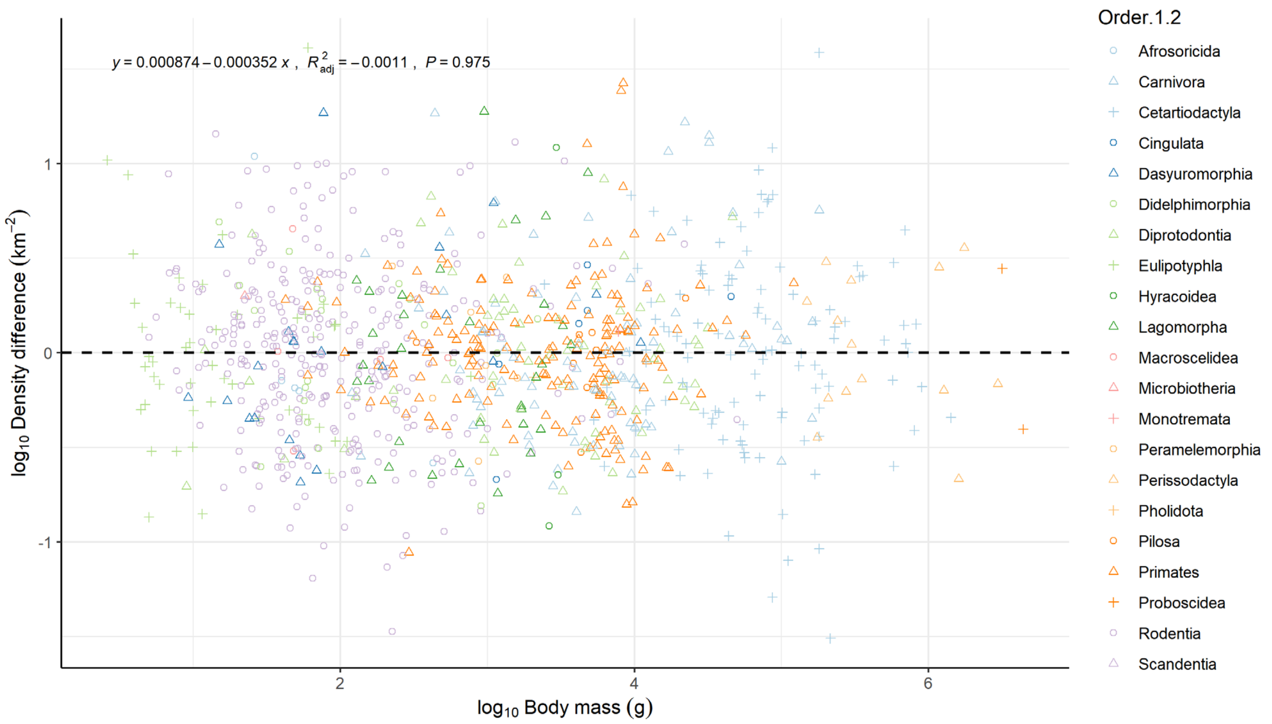


**Fig 4:** Overview of the difference in imputed values compared to the empirical estimates from PanTHERIA across body mass. Positive are model overestimates, negative are model underestimates. Color and shape indicate taxonomic order. The dashed line shows the 0, and the solid line indicates a linear fit. The plot shows a spread of values of the imputed densities about a factor 10 around the empirical values.

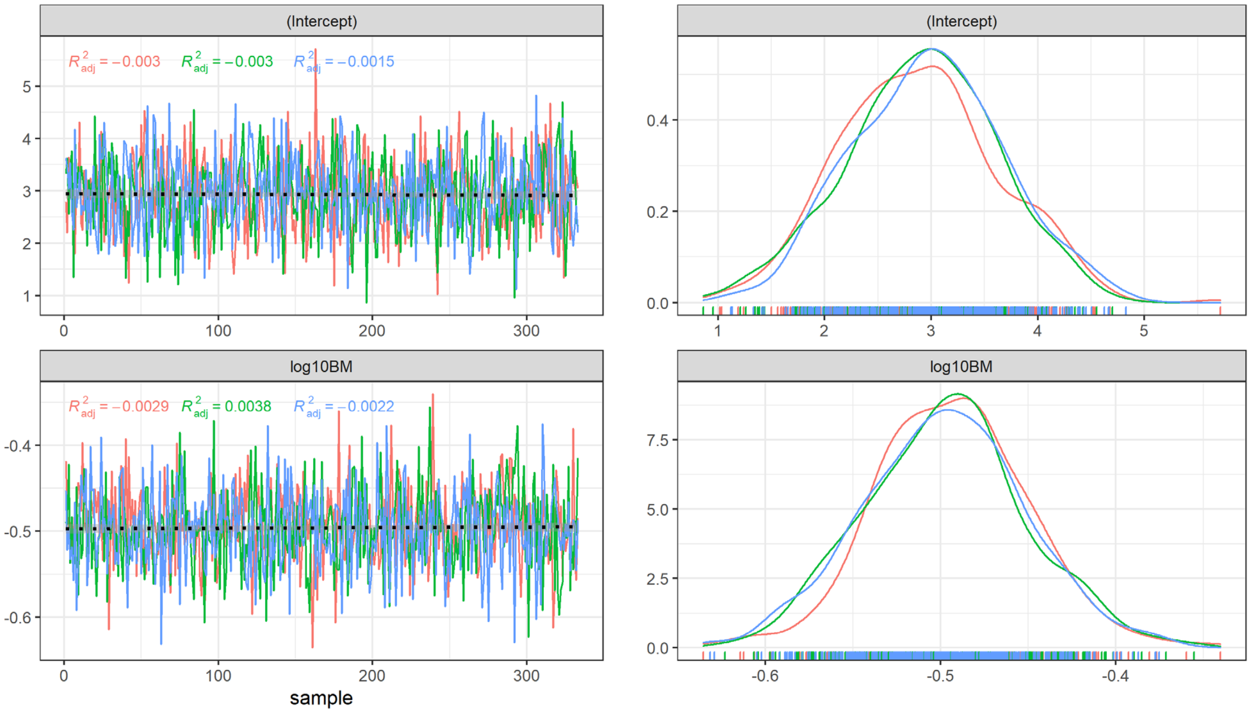


**Fig. 5:** Sample chains for main effects in the MCMCGLMM with Density as a function of body mass on a log-log scale. 3 chains are shown for testing purposes. The total set of 1000 trees was finally run and only sampled 3 times each.

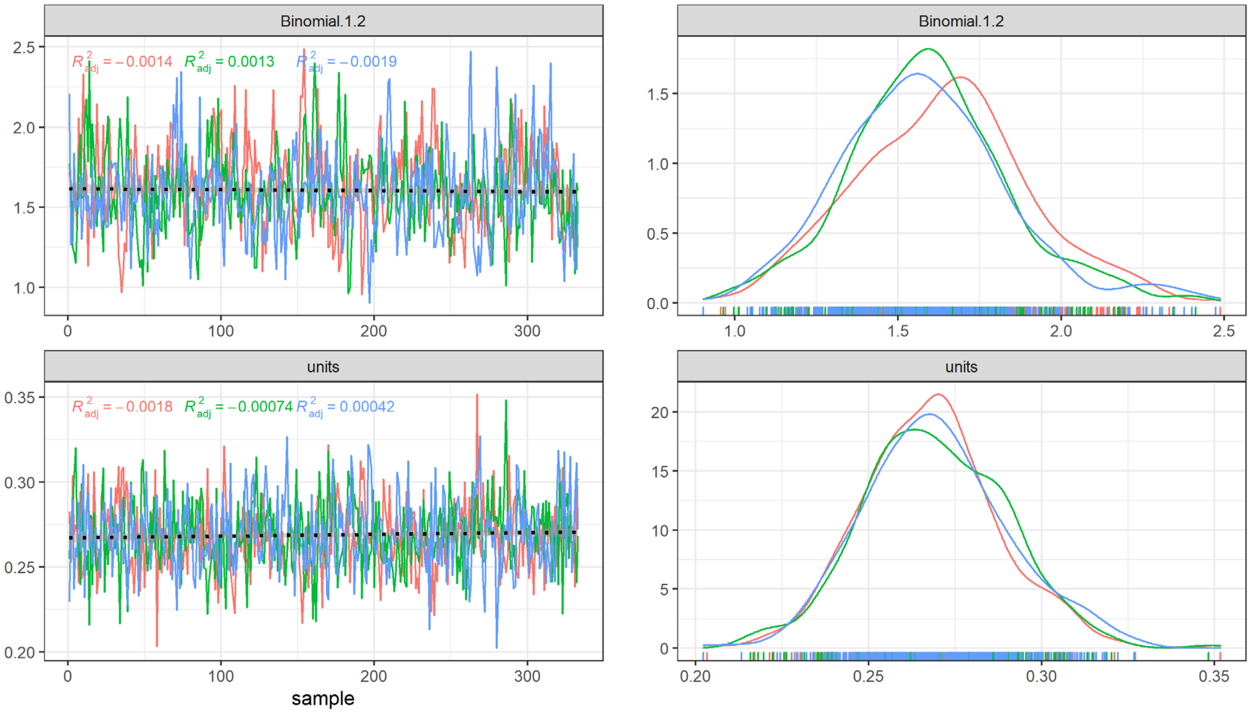


**Fig. 6:** Sample chains for random effects in the MCMCGLMM with Density as a function of body mass on a log-log scale. With a phylogeny as a random effect. 3 chains are shown for testing purposes. The total set of 1000 trees was finally run and only sampled 3 times each.

### References


Corporation, M., & Weston, S. (2017). doSNOW: Foreach Parallel Adaptor for the “snow” Package. Retrieved from https://cran.r-project.org/package=doSNOW

Faurby, S., Davis, M., Pedersen, R. Ø., Schowanek, S. D., Antonelli, A., & Svenning, J.-C. (2018). PHYLACINE 1.2: The Phylogenetic Atlas of Mammal Macroecology. *Ecology*, *99*(11), 2626–2626. https://doi.org/10.1002/ecy.2443

Faurby, S., Pedersen, R. Ø., Davis, M., Schowanek, S. D., Jarvie, S., Antonelli, A., & Svenning, J.-C. (2020). PHYLACINE 1.2.1: An update to the Phylogenetic Atlas of Mammal Macroecology. https://doi.org/10.5281/zenodo.3690867

Faurby, S., & Svenning, J.-C. (2015). A species-level phylogeny of all extant and late Quaternary extinct mammals using a novel heuristic-hierarchical Bayesian approach. *Molecular Phylogenetics and Evolution*, *84*, 14–26. https://doi.org/10.1016/j.ympev.2014.11.001

Faurby, S., & Svenning, J.-C. (2016). Resurrection of the Island Rule: Human-Driven Extinctions Have Obscured a Basic Evolutionary Pattern. *The American Naturalist*, *187*(6), 812–820. https://doi.org/10.1086/686268

Hadfield, J. (2012). MCMCglmm course notes. *See Http://Cran. r-Project. Org/Web/Packages/MCMCglmm/Vignettes/CourseNotes. Pdf*.

Hadfield, J. D. (2010). MCMC Methods for Multi-Response Generalized Linear Mixed Models: The {MCMCglmm} {R} Package. *Journal of Statistical Software*, *33*(2), 1–22. Retrieved from http://www.jstatsoft.org/v33/i02/

Jones, K. E., Bielby, J., Cardillo, M., Fritz, S. A., O’Dell, J., Orme, C. D. L., … Purvis, A. (2009). PanTHERIA: a species-level database of life history, ecology, and geography of extant and recently extinct mammals. *Ecology*, *90*(9), 2648–2648. https://doi.org/10.1890/08-1494.1

Paradis, E., Claude, J., & Strimmer, K. (2004). A{PE}: analyses of phylogenetics and evolution in {R} language. *Bioinformatics*, *20*, 289–290. Retrieved from https://cran.r-project.org/package=ape

R Core Team. (2017). R: A Language and Environment for Statistical Computing. Retrieved from http://www.r-project.org/

Wickham, H. (2017). tidyverse: Easily Install and Load the “Tidyverse.” Retrieved from https://cran.r-project.org/package=tidyverse
