## Appendix S2 for "Extinctions have strongly reduced the mammalian consumption of primary productivity"

### Metabolic rates

In the following we provide our method for imputing both basal and field metabolic rates for all terrestrial mammals, and provide a full summarised dataset of the results. We used both basal and field metabolic rates, since they are usually linked and provide a larger and more diverse dataset to work with across all mammals. Data provided as either field or basal could help inform the other in a model.

#### Data

We compiled metabolic rate estimates for specific body masses of individuals for both basal and field metabolic rates (Table S1b). We collected data from several published sources, filtering for duplicated data, and expanding from primary sources in phylogenetically scarce areas, as well as for large animals where data also was scarce. For taxonomy we followed the PHYLACINE database (v. 1.2.1) and their synonymy tables (Faurby et al., 2018, 2020). Further taxonomic resolution was done manually and is provided in the supplementary table (Table S1b). Data for basal metabolic rate not provided as energy but in quantities of $O_2$ were transformed using the following equation (adapted from Lusk, 1924) assuming respiration quotient (RQ) of 0.8.

$$[kcal] = (3.815 + 1.2321 * RQ) * [L\ O_2] = 4.8007 * [L\ O_2]$$

For phylogenetic imputation we used a posterior distribution of 1000 equally likely fully resolved phylogenies for all mammals provided with the PHYLACINE database (Faurby et al., 2018; Faurby & Svenning, 2015). Body mass for imputation of species specific metabolic rates was also acquired from PHYLACINE and underlying sources (Faurby et al., 2018; Faurby & Svenning, 2016; Smith et al., 2003).

The dataset was filtered to species classified as non-marine, non-flying, terrestrial species (i.e. excluding bats, whales, pinnipeds and sea cows, polar bears (*Ursus maritimus*) and two marine otters (*Enhydra lutris*, *Lontra felina*). The final dataset consisted of 591 species spanning 22 orders and 91 families, well distributed across the phylogeny and mass range (Fig 2) – though sparse in data for body mass > 100 kg, and missing > 1000 kg. Basal metabolic rate data was for 552 species with 753 data points and field metabolic rate data was for 115 species with 461 data points.

#### Imputation

We used the 'MCMCglmm' package v. 2.29 (J. D. Hadfield, 2010) in the R language v. 3.6.2 (R Core Team, 2017) for imputation, where we used relatively uninformative priors as recommended (J. Hadfield, 2012), G(V = 1, nu = 0.002) and R(V = 1, nu = 0.002). The model was specified as $\log_{10}$ *Density* as a function of $\log_{10}$ *Body mass* with the phylogeny as random effect. We found a burnin of 1000 with a thinning factor of 75 to be sufficient, which ran for 25 975 iterations on 3 chains giving 999 samples per species for one test tree. Effective sample size for intercept and slope was found to be 308 and 333 per chain, and 211 for the G-structure on phylogeny and 290 for the R-structure (Fig 7 and 8), Gelman diagnostics value was found < 1.02 for all variables. For the full dataset of trees only 3 samples per chain was necessary to give an overall impression of the value distribution across the 1000 trees. Across a posterior distribution of 1000 equally likely phylogenetic trees from PHYLACINE, giving 9000 records per species.

The following packages was used: 'doSNOW' v. 1.0.18 (Corporation & Weston, 2017) for parallel processing, 'ape' v. 5.3 (Paradis, Claude, & Strimmer, 2004) for handling phylogenies, and 'tidyverse' v. 1.3.0 (Wickham, 2017) for general data wrangling and plotting.

#### Results

The final results were summarised in two ways. The final dataset with 9000 samples for each species density was used throughout our calculations, and summarised everything with mean, median, and highest posterior distribution 95% credibility interval in Table S3b. We here provide only the estimated values with their uncertainties and ignore the original empirical values for a uniform dataset (Fig 1-2). Estimated densities follows the empirical values nicely and are relatively unbiased across the dataset (Fig 2-3).

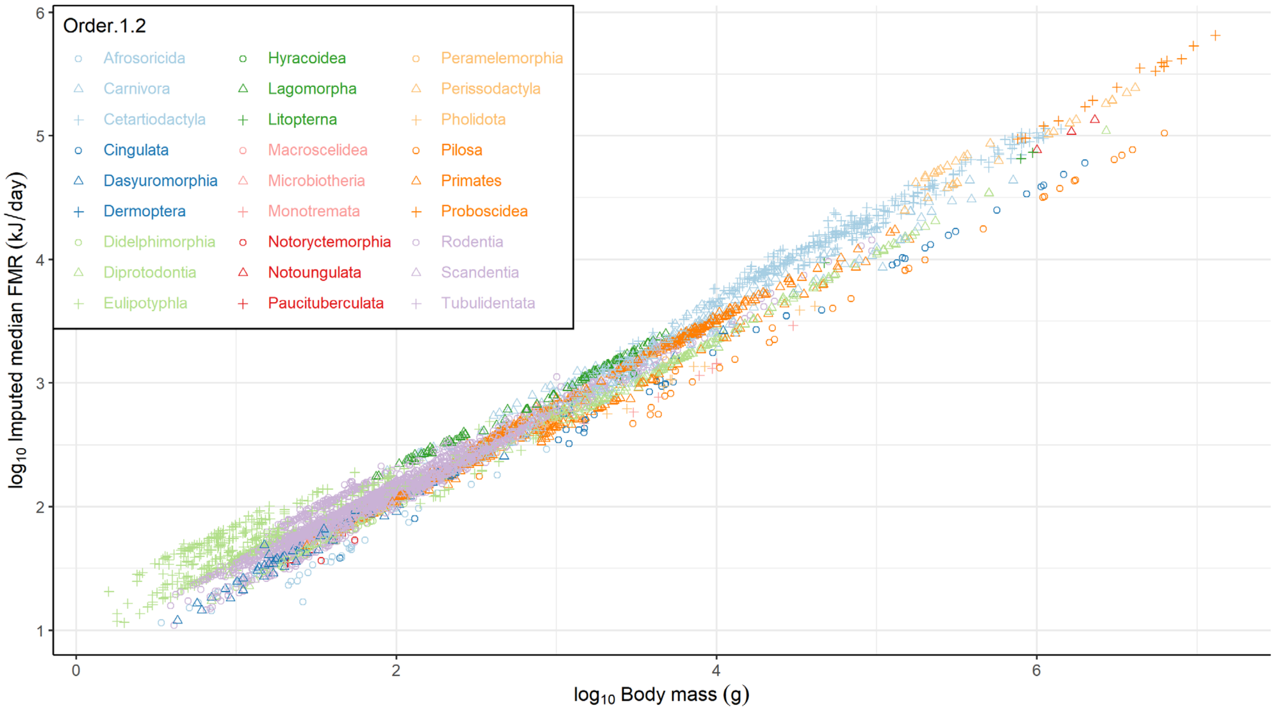


**Fig 1:** Median field metabolic rate estimates for all terrestrial mammals plotted along body size on a log10-log10 scale. The 29 orders of mammals are coded by a unique combination of symbol and colour to distinguish them as much as possible.

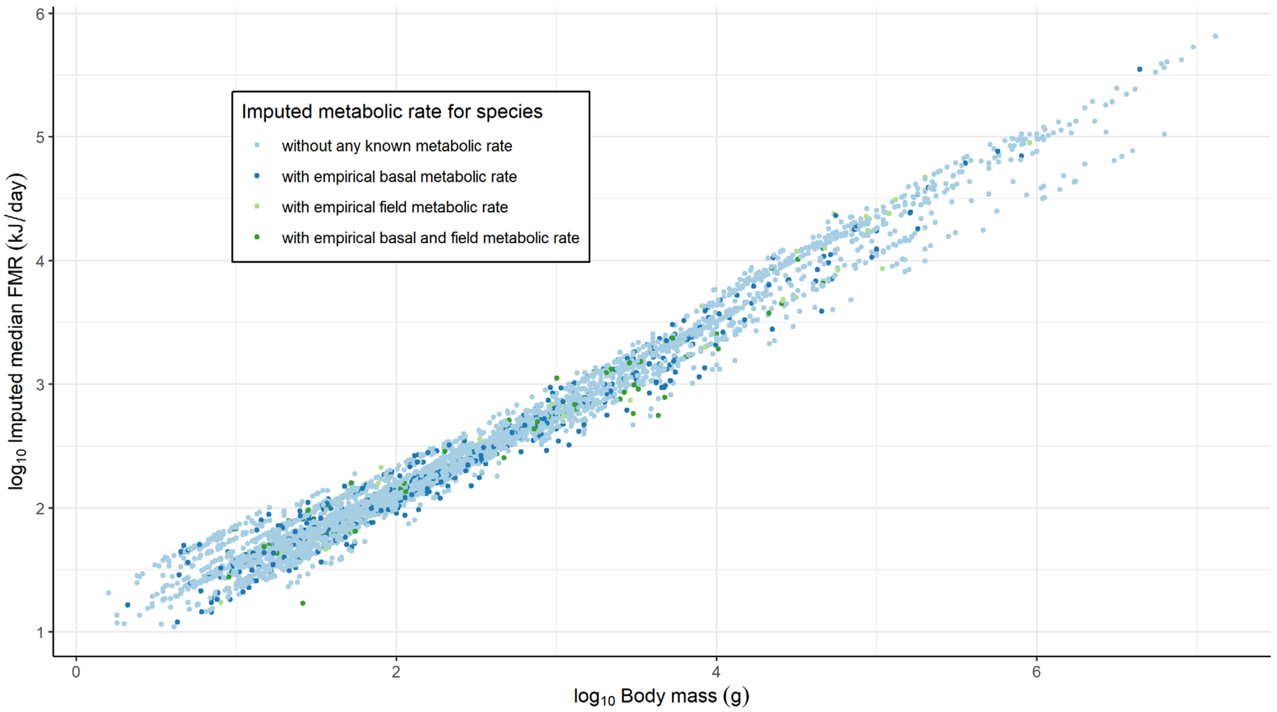


**Fig 2:** Resulting mean imputed field metabolic rates for all terrestrial mammals. All values are imputed and colours indicate whether we have underlying data for the species. For underlying data see Fig. 4.

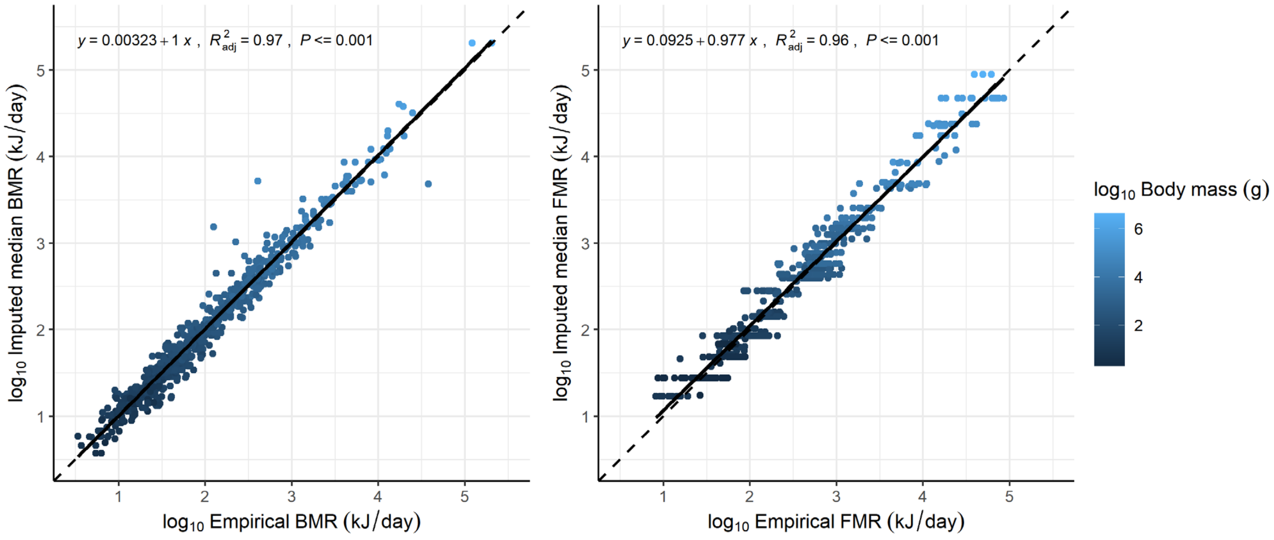


**Fig 3:** Overview of the imputed values compared to the empirical estimates. Color brightness shows body mass, the dashed line shows the 1:1 line, and the solid line indicates a linear fit. The plot shows a spread of values of the imputed densities about a factor 5 around the empirical values.

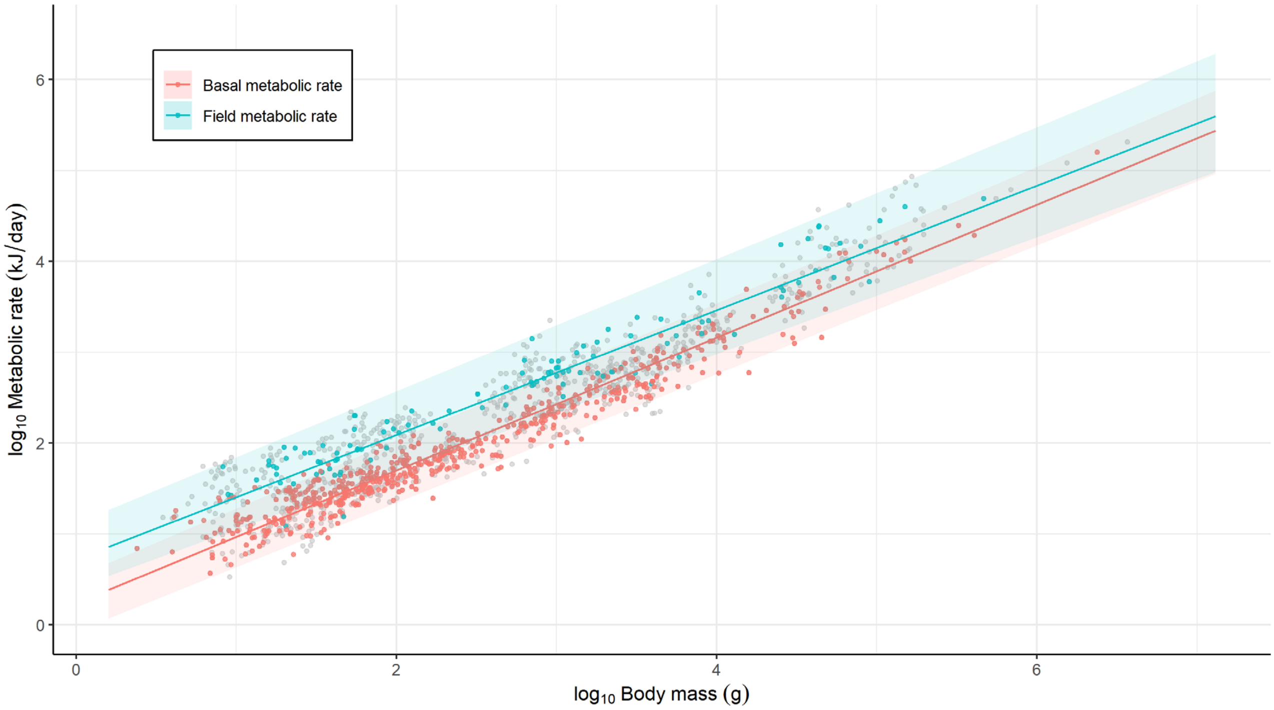


**Fig 4:** Underlying data for metabolic rate with fitted model on top. Grey point indicate all actual data values, while coloured points indicate either species empirical value, or mean species empirical values if more were available. Regression lines indicate overall fit of model with average phylogenetic random effect. For actual imputed estimates see Fig 5.

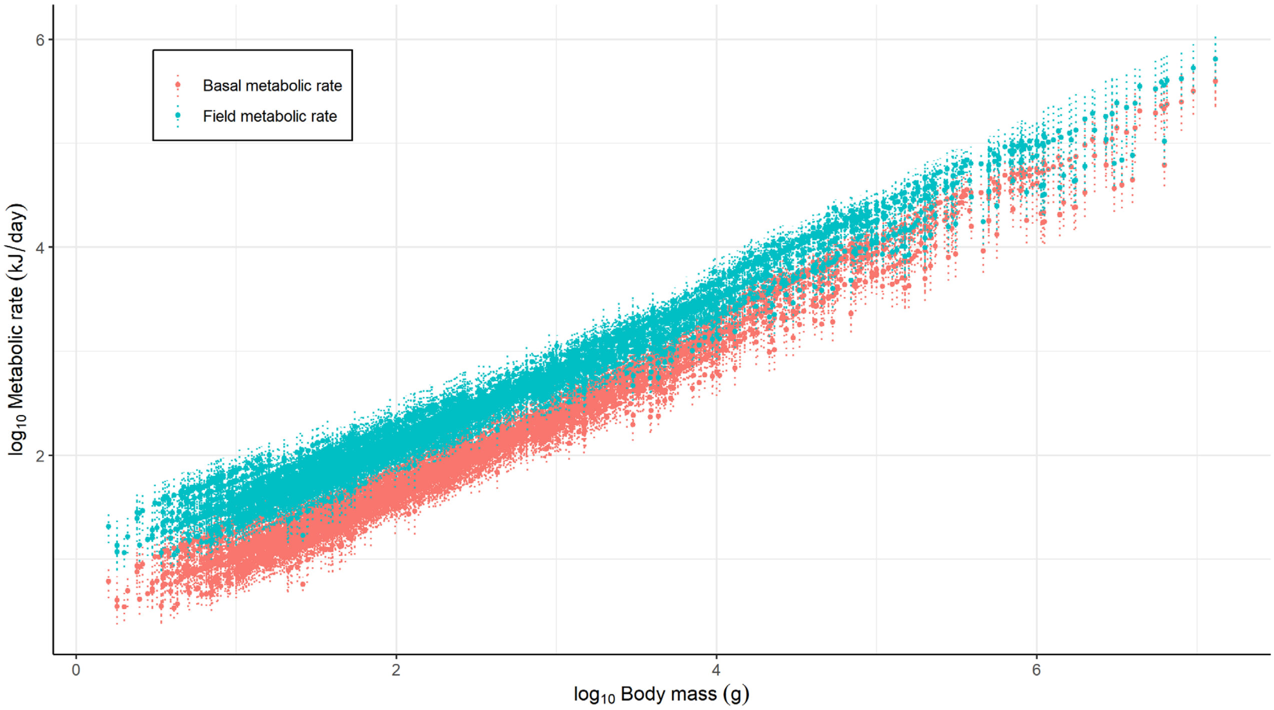


**Fig. 5:** Imputed basal and field metabolic rates for all terrestrial mammals living through the last 130 ka. Lines indicate 95% highest posterior density interval.

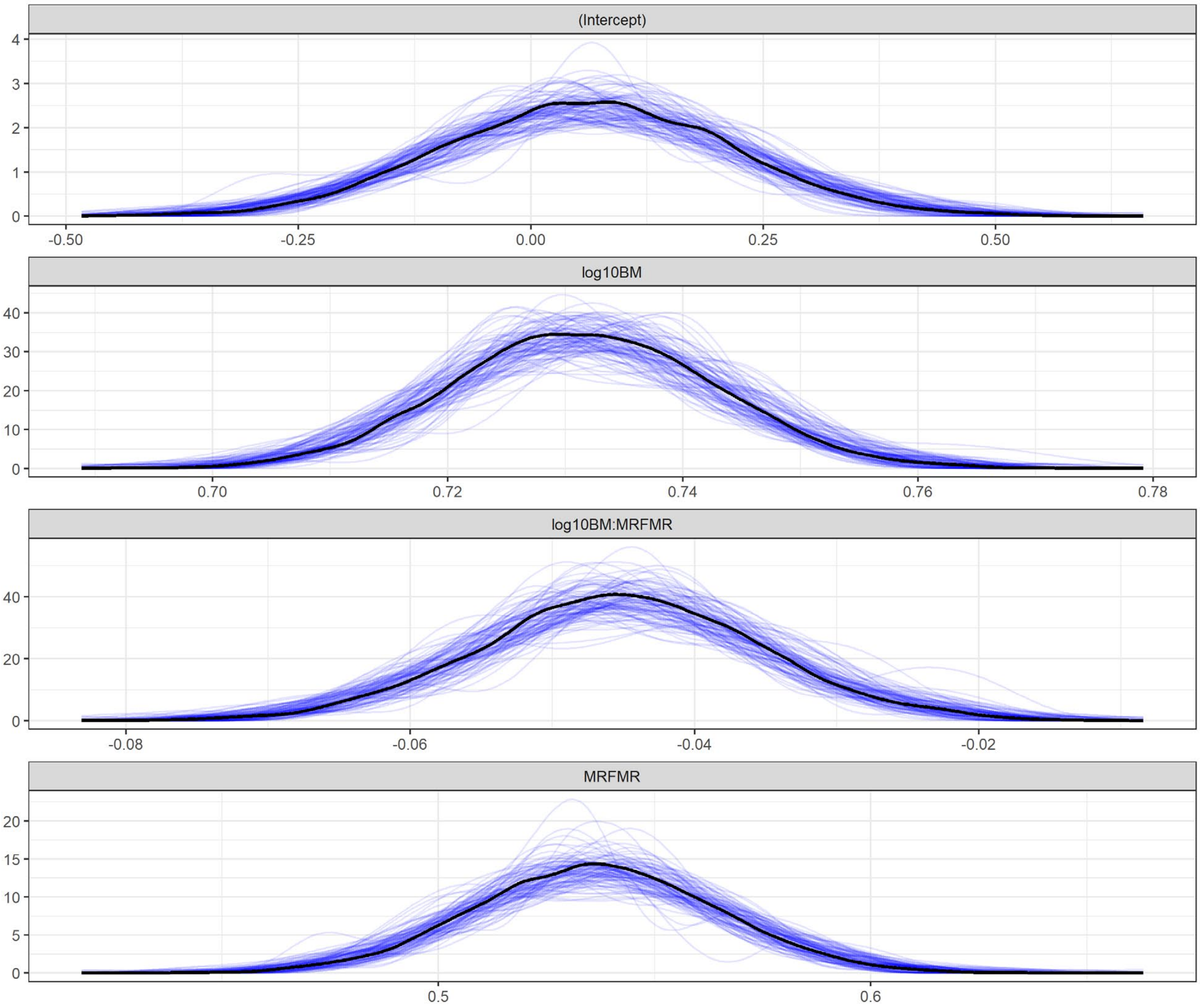


**Fig. 6:** MCMC-GLMM results. The overall density distribution of results are plotted black. The blue lines are results sampled per 10 trees. The results in the second row shows that metabolic rate on a log-log scale is related to body mass right between the two prevailing hypothesis the 2/3 scaling and 3/4 scaling. Row 4 shows the offset between BMR and FMR to be around 10^0.53 = 3.4 (95% CI: 3.02-3.90), slightly higher than the usually stated 3x. See Table 1 for details.

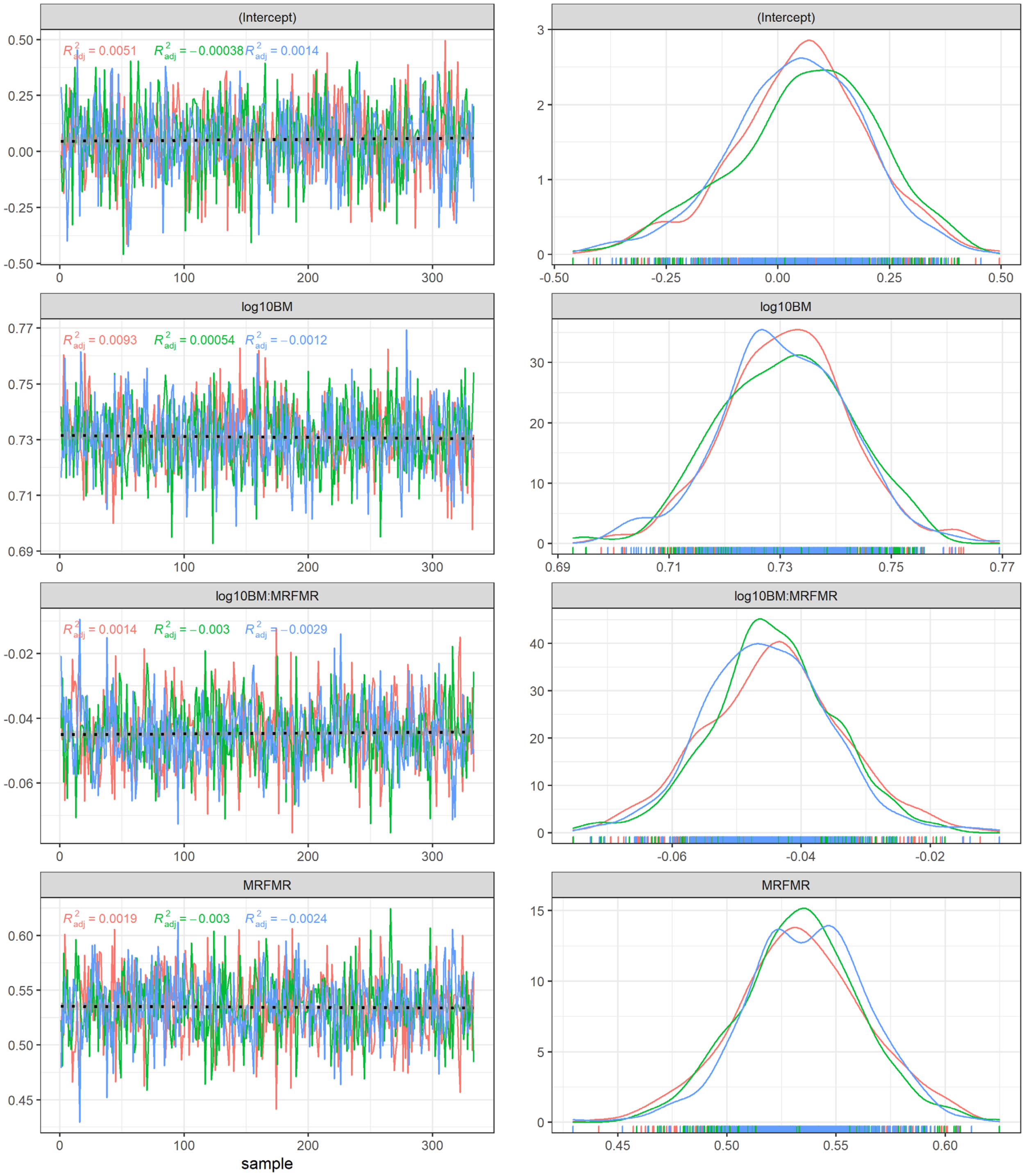


**Fig. 7:** Sample chains for main effects in the MCMCGLMM with Density as a function of body mass on a log-log scale. 3 chains are shown for testing purposes. The total set of 1000 trees was finally run and only sampled 3 times each.

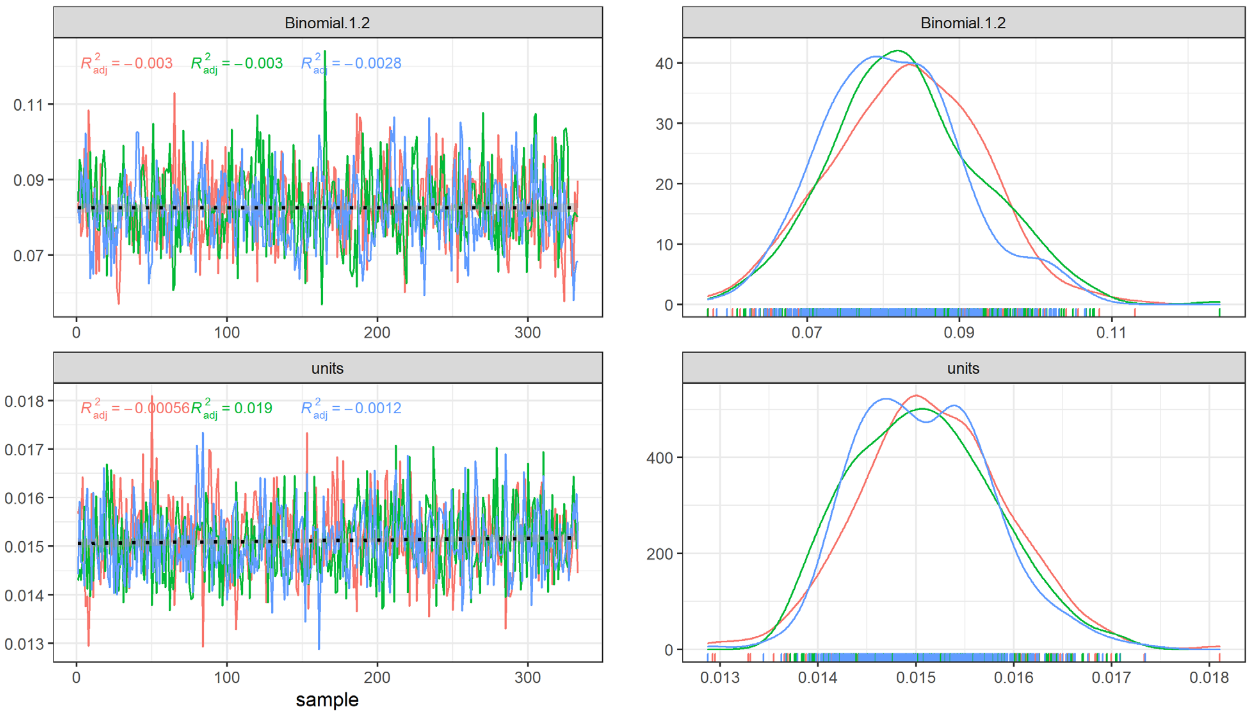


**Fig. 8:** Sample chains for random effects in the MCMCGLMM with Density as a function of body mass on a log-log scale. With a phylogeny as a random effect. 3 chains are shown for testing purposes. The total set of 1000 trees was finally run and only sampled 3 times each.

**Table 1:** Shows the median estimates and lower and upper HPD 95% confidence intervals.

| | median estimate | lower 95%-HPD interval | upper 95%-HPD interval |
|---|---|---|---|
| (Intercept) (Basal metabolic rate) | 0.062 | -0.25 | 0.35 |
| log10 Body mass (Basal metabolic rate) | 0.73 | 0.71 | 0.75 |
| Intercept offset (Field metabolic rate) | 0.54 | 0.48 | 0.59 |
| Slope change (Field metabolic rate) | -0.045 | -0.065 | -0.026 |
| Mean random effect | 0.18 | -0.12 | 0.47 |

### References

Corporation, M., & Weston, S. (2017). doSNOW: Foreach Parallel Adaptor for the “snow” Package. Retrieved from https://cran.r-project.org/package=doSNOW

Faurby, S., Davis, M., Pedersen, R. Ø., Schowanek, S. D., Antonelli, A., & Svenning, J.-C. (2018). PHYLACINE 1.2: The Phylogenetic Atlas of Mammal Macroecology. *Ecology*, *99*(11), 2626–2626. https://doi.org/10.1002/ecy.2443

Faurby, S., Pedersen, R. Ø., Davis, M., Schowanek, S. D., Jarvie, S., Antonelli, A., & Svenning, J.-C. (2020). PHYLACINE 1.2.1: An update to the Phylogenetic Atlas of Mammal Macroecology. https://doi.org/10.5281/zenodo.3690867

Faurby, S., & Svenning, J.-C. (2015). A species-level phylogeny of all extant and late Quaternary extinct mammals using a novel heuristic-hierarchical Bayesian approach. *Molecular Phylogenetics and Evolution*, *84*, 14–26. https://doi.org/10.1016/j.ympev.2014.11.001

Faurby, S., & Svenning, J.-C. (2016). Resurrection of the Island Rule: Human-Driven Extinctions Have Obscured a Basic Evolutionary Pattern. *The American Naturalist*, *187*(6), 812–820. https://doi.org/10.1086/686268

Hadfield, J. (2012). MCMCglmm course notes. *See Http://Cran. r-Project. Org/Web/Packages/MCMCglmm/Vignettes/CourseNotes. Pdf*.

Hadfield, J. D. (2010). MCMC Methods for Multi-Response Generalized Linear Mixed Models: The {MCMCglmm} {R} Package. *Journal of Statistical Software*, *33*(2), 1–22. Retrieved from http://www.jstatsoft.org/v33/i02/

Lusk, G. (1924). Animal calorimetry. Twenty-fourth paper. Analysis of the oxidation of mixtures of carbohydrate and fat. A correction. *Journal of Biological Chemistry*, *59*(1), 41–42.

Paradis, E., Claude, J., & Strimmer, K. (2004). A{PE}: analyses of phylogenetics and evolution in {R} language. *Bioinformatics*, *20*, 289–290. Retrieved from https://cran.r-project.org/package=ape

R Core Team. (2017). R: A Language and Environment for Statistical Computing. Retrieved from http://www.r-project.org/

Smith, F. A., Lyons, S. K., Ernest, S. K. M., Jones, K. E., Kaufman, D. M., Dayan, T., … Haskell, J. P. (2003). Body mass of late quaternary mammals. *Ecology*, *84*(12), 3403–3403. https://doi.org/10.1890/02-9003

Wickham, H. (2017). tidyverse: Easily Install and Load the “Tidyverse.” Retrieved from https://cran.r-project.org/package=tidyverse
